# Microcephaly-associated WDR62 mutations hamper Golgi apparatus-to-spindle pole shuttling in human neural progenitors

**DOI:** 10.1101/2022.07.24.501306

**Authors:** Claudia Dell’Amico, Marilyn M. Angulo Salavarria, Yutaka Takeo, Ichiko Saotome, Maria Teresa Dell’Anno, Maura Galimberti, Enrica Pellegrino, Elena Cattaneo, Angeliki Louvi, Marco Onorati

## Abstract

WDR62 is a spindle pole-associated scaffold protein with pleiotropic functions during corticogenesis. Recessive mutations in WDR62 are associated with structural brain abnormalities and account for the second most common cause of autosomal recessive primary microcephaly (MCPH), indicating WDR62 as a critical hub for human brain development. Here, we investigated a C-terminal truncating mutation (D955AfsX112) in WDR62 using induced pluripotent stem cells (iPSCs) obtained from a patient with MCPH2. We generated neuroepithelial stem (NES) cells and cerebro-cortical progenitors and neurons from patient-derived and isogenic retro-mutated iPSC lines. We found that WDR62 dysfunction resulted in impaired cell cycle progression and alterations of the neurogenic trajectories of iPSC neuroderivatives. Moreover, we report WDR62 localization at the Golgi apparatus during interphase, both in human neural progenitors in vitro and in human fetal brain tissue. WDR62 shuttling from the Golgi apparatus to spindle poles is dynamic and microtubule-dependent. Impairment of WDR62 function and localization results in severe neurodevelopmental abnormalities, thus delineating new mechanisms in MCPH etiology.

## Introduction

The development of the human brain is a sophisticated process that extends over several decades, during which a plethora of distinct cell types are generated and assembled into functionally distinct circuits and regions (Lui et al., 2011; Silbereis et al., 2016). The precise orchestration and coordination of neural stem/progenitor cell proliferation and differentiation are pivotal for the formation and function of the central nervous system (CNS). Deviations from normal neurodevelopmental processes, e.g., cell cycle progression, symmetric versus asymmetric cell divisions, lineage commitment, as well as neuronal migration and differentiation, ultimately affect the structure and function of the CNS and can lead to neurological or psychiatric disorders (M. Li et al., 2018).

Mutations in genes involved in the regulation of mitotic progression in neural progenitor cells (NPCs) have been identified in several types of MCPH (Phan & Holland, 2021; Thornton & Woods, 2009), a genetically heterogeneous disorder characterized by occipito-frontal circumference 3-4 standard deviations below the mean of ethnically-, age-, and sex-matched controls (Woods & Parker, 2013). Intriguingly, many of the genes associated with MCPH encode centrosomal or pericentriolar proteins, while others participate in mitotic spindle organization and mitotic progression (Jayaraman et al., 2018; Phan & Holland, 2021).

Recessive mutations in WDR62, located on chromosome 19, are responsible for the second most frequent form of MCPH (Bilgüvar et al., 2010; Nicholas et al., 2010; Yu et al., 2010). WDR62 encodes a WD-domain containing protein, member of an ancient large family of proteins involved in coordinating multiprotein assemblies, with the WD repeats serving as a scaffold for protein interactions (Shohayeb et al., 2018). WD-repeat proteins have pleiotropic functions, ranging from signal transduction and transcriptional regulation to cell cycle control and apoptosis (Jain & Pandey, 2018; D. Li & Roberts, 2001; Smith et al., 1999; Stirnimann et al., 2010). WDR62 is a spindle pole-associated protein involved in mitotic progression and NPC maintenance (Sgourdou et al., 2017). In addition, it has a pivotal role in the assembly and disassembly of the primary cilium (Shohayeb et al., 2020; Zhang et al., 2019), an antenna-like structure able to sense and convey extracellular cues guiding NPC proliferation versus differentiation (Wilsch-Bräuninger & Huttner, 2021). The subcellular localization of WDR62 is cell-cycle dependent and regulated by its N and/or C-terminal domains and microtubule association (Bogoyevitch et al., 2012). Notably, in interphase, WDR62 is enriched in the centrosome via stepwise hierarchical interactions with other MCPH-associated proteins, including CDK5RAP2 at the top and CEP63 at the bottom of the cascade (Kodani et al., 2015; O’Neill et al., 2022). This complex is then stabilized by centriolar satellite proteins and is required for proper centriole duplication (Kodani et al., 2015).

We previously reported a homozygous truncating mutation (D955AfsX112) in WDR62 in patients with microcephaly and structural brain abnormalities (Sgourdou et al., 2017). In the present work, we directly investigated the effects of this mutation on NPC proliferation and neurogenic potential, taking advantage of patient-derived induced pluripotent stem cells (iPSCs) and isogenic lines generated via CRISPR/Cas9 gene editing (Dell’ Amico et al., 2021). From WDR62 and Isogenic iPSCs we derived neuroepithelial stem (NES) cells, the founders of cerebro-cortical cellular complexity (Baggiani et al., 2020; Onorati et al., 2016), as well as terminally differentiated cerebro-cortical neurons, through a directed differentiation protocol (Chambers et al., 2009; Shi et al., 2012).

Our results suggest that the mutation under study impairs cell cycle progression, leading to lineage specification defects in iPSC-derived cerebrocortical neurons. We also found that WDR62 dynamically associates with the Golgi apparatus, both in NES cells and human fetal telencephalic tissue. During the interphase-to-mitosis transition, WDR62 shuttles from the Golgi apparatus to the spindle poles in a microtubule-dependent manner. Nocodazole treatment of Isogenic control NES cells mimics the effects of the genetic mutation, preventing WDR62 shuttling from the Golgi apparatus to the spindle poles during cell cycle progression. These findings, together with perturbations we observed in the timing of differentiation of cerebro-cortical neurons from mutant iPSCs, suggest novel potential mechanisms underlying WDR62 involvement in human neurodevelopment and MCPH etiology.

## Results

### The D955AfsX112 mutation produces a truncated WDR62 protein and disrupts spindle pole localization

We previously reported a consanguineous family with two affected siblings displaying global developmental delay (Sgourdou et al., 2017). The index case, a 13-year-old male, and his 6-year-old brother, presented to medical attention for global developmental delay, severe microcephaly, and dysmorphic facial traits. Neuroimaging revealed a plethora of developmental abnormalities including diffuse pachygyria, thickened cortex, hypoplastic corpus callosum, and metopic synostosis. Whole exome sequencing identified a homozygous 4bp deletion in exon 23 of WDR62, leading to a frameshift and a premature STOP codon (a.a. 1067), which resulted in a C-terminally truncated peptide (D955AfsX112) (Figure 1A). The mutation was confirmed to be homozygous in both affected subjects, and heterozygous in both parents by Sanger sequencing (Sgourdou et al., 2017). To investigate possible alterations of cortical development linked to the mutation, fibroblasts (obtained from skin biopsies from family members) were reprogrammed to iPSCs (“WDR62 iPSCs”) using episomal vectors, assessed for pluripotency, and subsequently differentiated to relevant neural populations, i.e., NES cells and mature cerebro-cortical neurons (Figure 1B and Figure 1-figure supplement 1) (Chambers et al., 2009). To obtain a gold-standard control for all subsequent analyses, we corrected the mutation through CRISPR/Cas9 gene editing technology in the WDR62 iPSC line, generating isogenic controls (hereafter “Iso iPSCs”). We verified sequence restoration in 5 clones by Sanger sequencing (Figure 1B); three randomly selected clones were verified for pluripotency (Figure 1-figure supplement 1) and further analyzed. We examined WDR62 expression in WDR62 iPSCs by confocal imaging and found that the mutation impairs spindle pole localization during mitosis, with mutant WDR62 appearing diffuse around the nucleus (Figure 1C). In contrast, in external control (CTRL) iPSCs, WDR62 was detected at the spindle poles, identified by PCNT and TUBA1A co-staining, (Figure 1C and Figure 1-figure supplement 2 A-C), as we reported previously (Sgourdou et al., 2017). The WDR62 signal was restored at the spindle poles in mitotic Iso iPSCs (Figure 1C). Quantification of WDR62 signal distribution through fluorescence intensity analysis revealed high and narrow fluorescence peaks in Iso iPSCs, similarly to CTRL iPSCs (Figure 1D); in contrast, no such peaks were detected in patient-derived iPSCs (Figure 1C, D).

**Figure 1:**
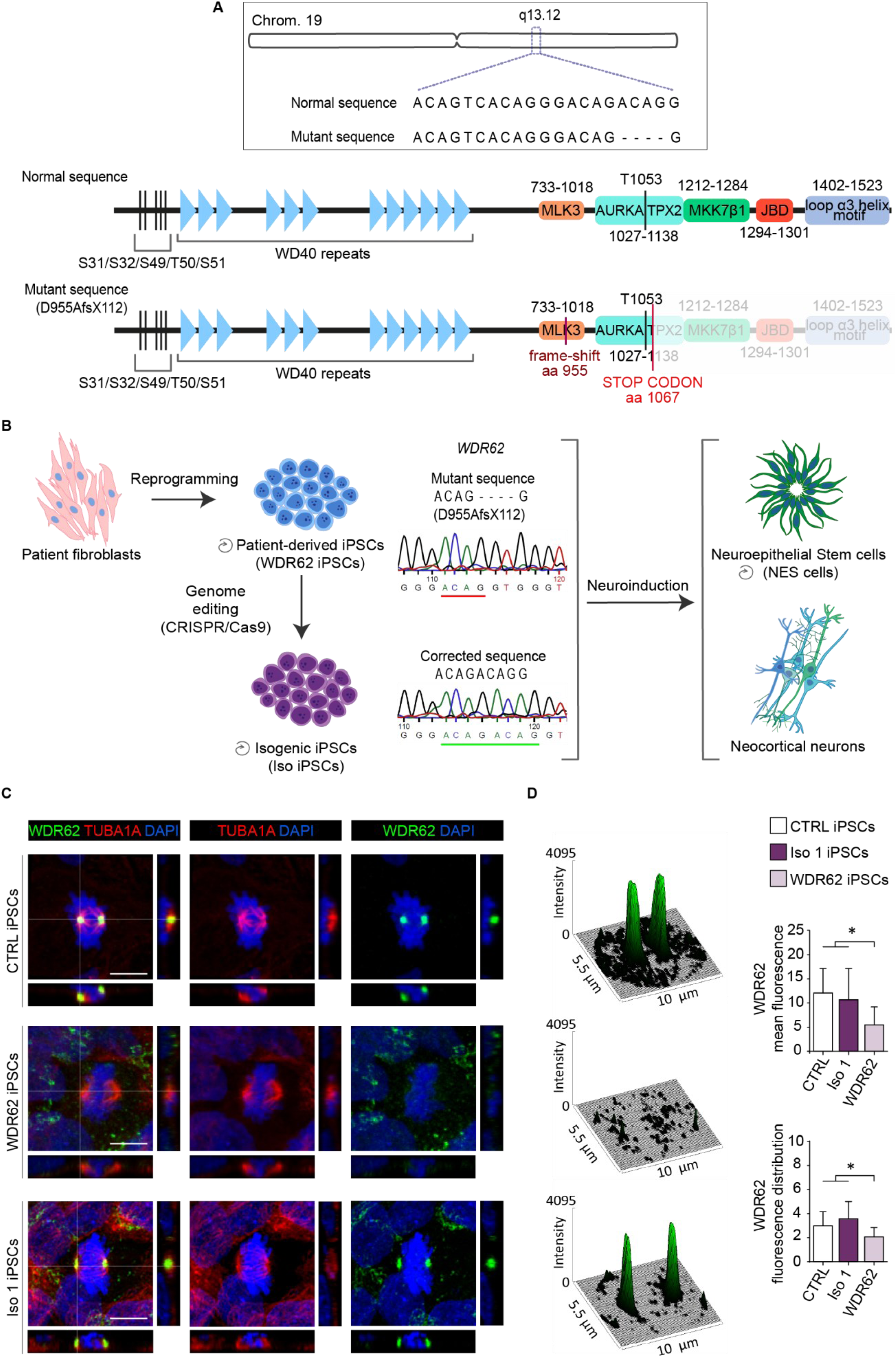
The D955AfsX112 mutation disrupts WDR62 spindle pole localization during mitosis. A) Schematic illustration of WDR62 gene structure on chromosome 19q13. The mutation is characterized by 4 bp (ACAG) deletion in exon 23 leading to a frameshift (D955AfsX112) and a C-terminally truncated protein. B) Scheme of experimental design. WDR62 iPSCs harboring the homozygous mutation and isogenic-corrected (Iso) iPSC lines were employed. Sanger sequencing of iPSCs shows the above-described mutation (top) and the CRISPR/Cas9 corrected sequence (bottom). A cerebro-cortical differentiation protocol was applied to WDR62 and Iso 1 and/or Iso 2 iPSCs to obtain relevant neural populations. C) Representative confocal images of WDR62 and alpha-tubulin (TUBA1A) expression in CTRL (external control), WDR62 (mutant) and Iso 1 iPSCs during mitosis. D) Surface plots and fluorescence intensity analysis show WDR62 signal distribution (fluorescence distribution and mean fluorescence, arbitrary unit – a.u.) during metaphase. The CTRL and Iso 1 iPSC lines show similar WDR62 signal distribution, which is decreased in the measured area in WDR62 iPSCs (N = 47, p-value < 0.05, Kruskal-Wallis test, post hoc Dunn’s test). Data are shown as mean ± SD. Scale bar = 10 μm.

### The D955AfsX112 WDR62 mutation impacts cell cycle progression in NES cells

To investigate the cellular and molecular consequences of the mutation in the context of neurodevelopment, we differentiated WDR62- and Iso iPSCs toward neural fate by applying a neocortical NES cell derivation protocol (Chambers et al., 2009) (Figure 2A). Briefly, after initial induction of dorsal forebrain neuroepithelial identity, based on Dual SMAD inhibitors and the WNT inhibitor XAV, we captured long-term self-renewing populations of iPS-derived NES (iPS-NES) cells by adding growth factors.

**Figure 2:**
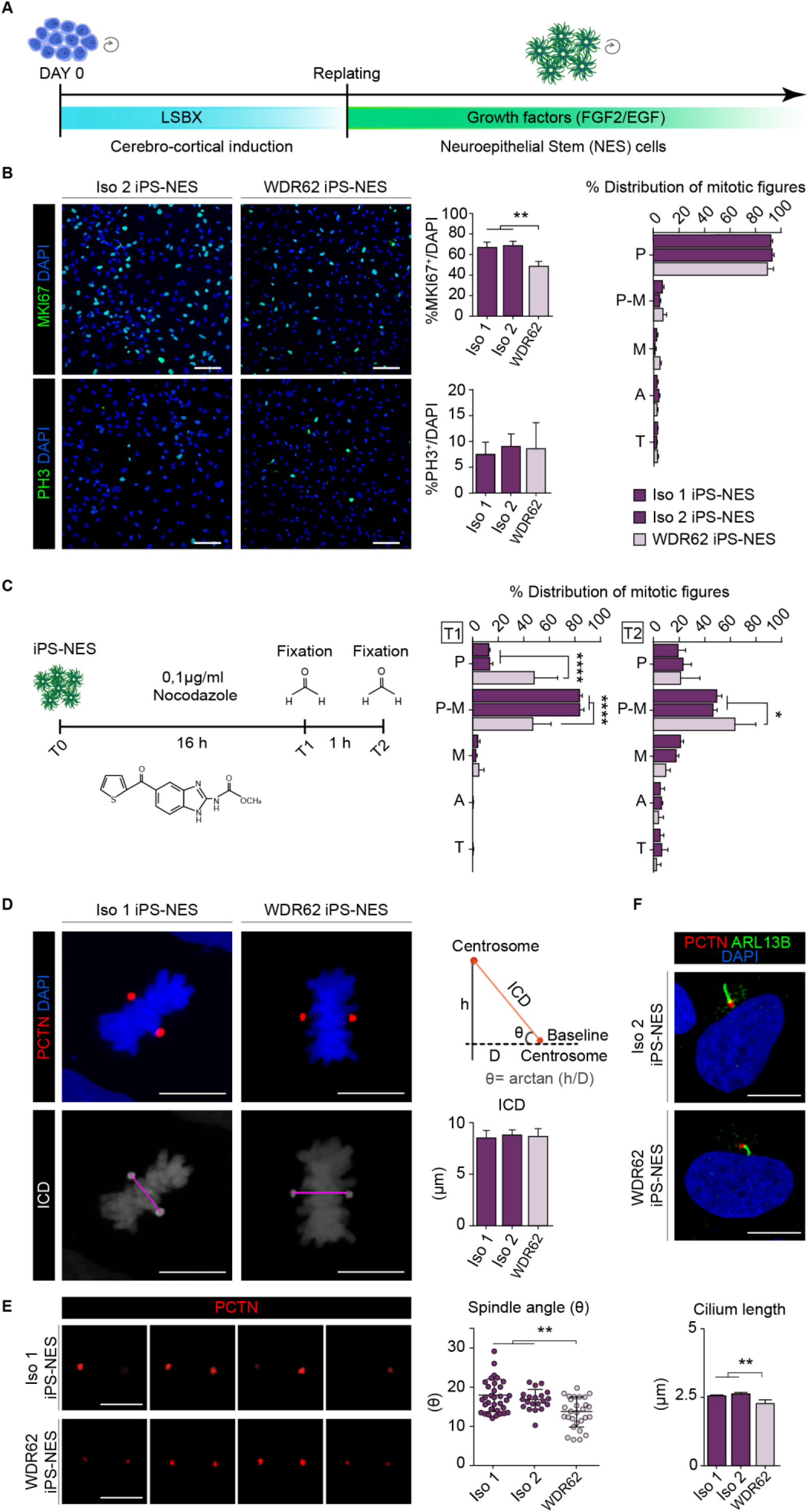
WDR62 iPS-NES cells exhibit defects in cell cycle regulation. A) Schematic illustration of NES cell derivation protocol. B) Representative confocal images of MKI67 and PH3 in Iso and WDR62 iPS-NES cell lines. Quantitative analysis for MKI67 and PH3 reveals a decrease in G_1_-M (MKI67^+^, N = 3315), but not in mitotically active cells (PH3^+^, N = 12684), p-value < 0.001 and p-value > 0.05, respectively. One-way ANOVA, post-hoc Tukey’s test. Mitotic figure distribution in unsynchronized cell cultures (global mitotic index) does not show significant differences between WDR62 and Iso iPS-NES cell lines (N = 12684, p-value < 0.05, two-way ANOVA, post hoc Dunnett’s test). P (prophase); P-M (prometaphase); M (metaphase); A (anaphase) and T (telophase). C) Scheme of synchronization experiment in iPS-NES cells. Immunofluorescence and quantitative analysis after 16 h of treatment (T1) show increased mitotic cell fraction in P-M. The percentage of synchronized P-M cells is higher in Iso iPS-NES than in WDR62 iPS-NES cultures. Fewer WDR62 iPS-NES cells proceed to M following nocodazole treatment and subsequent release in standard culture conditions (T2) compared to Iso iPS-NES cells (N = 8764, p-value < 0.0001, two-way ANOVA, post hoc Dunnett’s test and N = 9751, p-value < 0.05, two-way ANOVA, post hoc Dunnett’s test). D) Representative confocal images of PCTN and DAPI staining in Iso iPS-NES and WDR62 iPS-NES cells in M and example of inter-centrosome distance measurements (ICD) and spindle angle (θ). Quantitative analyses show no differences in ICD (N = 81, p-value > 0.05, one-way ANOVA, post hoc Sidak’s test). E) Confocal images of deconstructed Z-stacks of PCTN signal in iPS-NES cell lines in M. Spindle angle (θ) is smaller in WDR62 iPS-NES compared with Iso iPS-NES cells (N = 81, p-value < 0.01, one-way ANOVA, post hoc Dunnett’s test). F) Immunofluorescence analysis of PCTN and ARL13B in iPS-NES cells during interphase. Quantitative analysis shows shorter primary cilia in WDR62 iPS-NES cells (N = 850, p-value < 0.01, one-way ANOVA, post hoc Dunnett’s test). Data are shown as mean ± SD. Scale bar = 100 μm in B and 10 μm in D, E, F.

By consensus, MCPH is thought to be a consequence of neural progenitor depletion/premature differentiation (Jayaraman et al., 2018; Phan & Holland, 2021). For instance, in MCPH3 (caused by mutations in CDK5RAP2) (Lancaster et al., 2013), MCPH5 (ASPM) (R. Li et al., 2017), and MCPH6 (CENPJ) (Bond et al., 2005), neural progenitors at the Ventricular Zone (VZ)/Sub-Ventricular Zone (SVZ) lose their broad proliferative capacity impacting final brain architecture and size (Gabriel et al., 2020; Phan & Holland, 2021). For this reason, we sought to determine whether the WDR62 mutation might impact NES cell features. We first assessed WDR62 mRNA and protein production in WDR62 iPS-NES cells. We found no differences in the expression level of WDR62 by RT-qPCR between Iso and WDR62 iPS-NES cells (Figure 2-figure supplement 1A). We also detected by Western blot the WDR62 mutant protein (Figure 2-figure supplement 1B and Figure 2-figure supplement 1-source data 1).

We performed phenotypic analysis on normal versus WDR62 mutant NES cells. By staining with MKI67 (also known as Ki-67) and PH3, markers of proliferating cells, we found significantly fewer MKI67^+^ cells in WDR62 iPS-NES cultures (Figure 2B). Since MKI67 detects cells at all phases of the cell cycle except G_0_, this result suggests that WDR62 iPS-NES cells show a tendency for premature cell cycle exit. On the other hand, we did not find significant differences in the global mitotic index (PH3^+^/total cells) or mitotic figure distribution (i.e., cells at each mitotic phase, evaluated through PH3 staining), between WDR62, Iso 1 and Iso 2 iPS-NES cells in unsynchronized cultures (Figure 2B).

Aware that growth factor supplementation can propel NES cell proliferation, we further investigated cell cycle progression using additional methods. We induced cell cycle arrest through nocodazole - a microtubule depolymerizing agent (Figure 2C) - to synchronize iPS-NES cell cultures and evaluate their capability to reorganize cytoskeletal components, such as the mitotic spindle, and proceed through mitosis. As previously reported (Matsui et al., 2012; Yiangou et al., 2019), nocodazole treatment induces prometaphase arrest synchronizing the cultures after 16 h (T1). Following nocodazole removal and 1 h release (T2), cells re-enter the cell cycle. Our analysis revealed that, compared to Iso 1 and Iso 2 iPS-NES cells, the percentage of WDR62 iPS-NES cells in prometaphase was reduced at T1, but increased at T2 (Figure 2C). Iso iPS-NES cells responded to the treatment as expected, with more than 80% of cells arrested in prometaphase at T1 re-entering the cell cycle at T2. Therefore, our data confirm that WDR62 iPS-NES cells, after an insult affecting cell cycle progression, are delayed compared to Iso counterparts.

Given previously described functions of WDR62 in regulating mitotic spindle stability and mitotic spindle angle (Bogoyevitch et al., 2012; Guerreiro et al., 2021; Miyamoto et al., 2017), and primary cilium assembly/disassembly (Shohayeb et al., 2020; Zhang et al., 2019), we evaluated inter-centrosomal distance (ICD) and spindle pole angle, two indicators of mitotic spindle formation and integrity. ICD, measured as edge-to-edge pericentrin (PCTN) signal, which localizes in close proximity to the centrosome, was not altered in WDR62 iPS-NES cells (Figure 2D). On the contrary, the spindle angle was smaller in WDR62 iPS-NES cells compared to Iso 1 and Iso 2 iPS-NES cells (p < 0.05) (Figure 2E), thus suggesting mitotic spindle abnormalities.

WDR62, together with CEP170 and KIF2A, has been proposed to promote primary cilia assembly (Zhang et al., 2019). This prompted us to test primary cilia length in WDR62 iPS-NES cells through staining with PCTN, which is localized at the basal body during interphase (Bettencourt-Dias et al., 2011) and ARL13B, a marker of primary cilia (Figure 2F). Primary cilia appeared significantly shortened in WDR62 iPS-NES cells, compared to Iso counterparts (Figure 2F). Taken together, these results suggest a general disruption of mitotic progression in WDR62 iPS-NES cells, most likely due to spindle defects and primary cilium impairment.

### The D955AfsX112 mutation disrupts WDR62 Golgi-to-spindle pole shuttling

In addition to its transient association with the spindle poles during mitosis in iPSCs, we detected WDR62 as a polarized perinuclear signal during interphase, which appeared to be localized around the centrosome (marked by PCTN) in G_1_-S in iPSCs (Figure 3-figure supplement 1). Because the Golgi apparatus is known to be in close physical proximity to the centrosome (Sütterlin & Colanzi, 2010), we examined a potential association of WDR62 with the Golgi apparatus in Iso and WDR62 iPS-NES cells by immunofluorescence.

In Iso iPS-NES cells during mitosis, the WDR62 signal was centered, as expected, on discrete regions corresponding to the spindle poles (TUBA1A-positive) (Figure 3A). On the other hand, the Golgi signal (GOLGA1/Golgin 97-positive) was diffuse and scattered around the chromatin, representing the clusters forming during mitosis that are eventually partitioned between daughter cells (Ayala et al., 2020). In contrast, in WDR62 iPS-NES during mitosis, the WDR62 signal was diffuse and overlapped with GOLGA1 in more than 60% of cells analyzed (Figure 3B and Figure 3-figure supplement 2A). Despite its divergent behavior in mitotic WDR62 and Iso iPS-NES cells, WDR62 localized to the Golgi apparatus during interphase in iPS-NES cells of both genotypes (Figure 3C and Figure 3-figure supplement 2B). Furthermore, analysis of fluorescence signal intensity suggested that full-length and mutant WDR62 localize to the Golgi apparatus at comparable amounts, potentially excluding a non-physiological accumulation of the mutant protein (Figure 3C).

**Figure 3:**
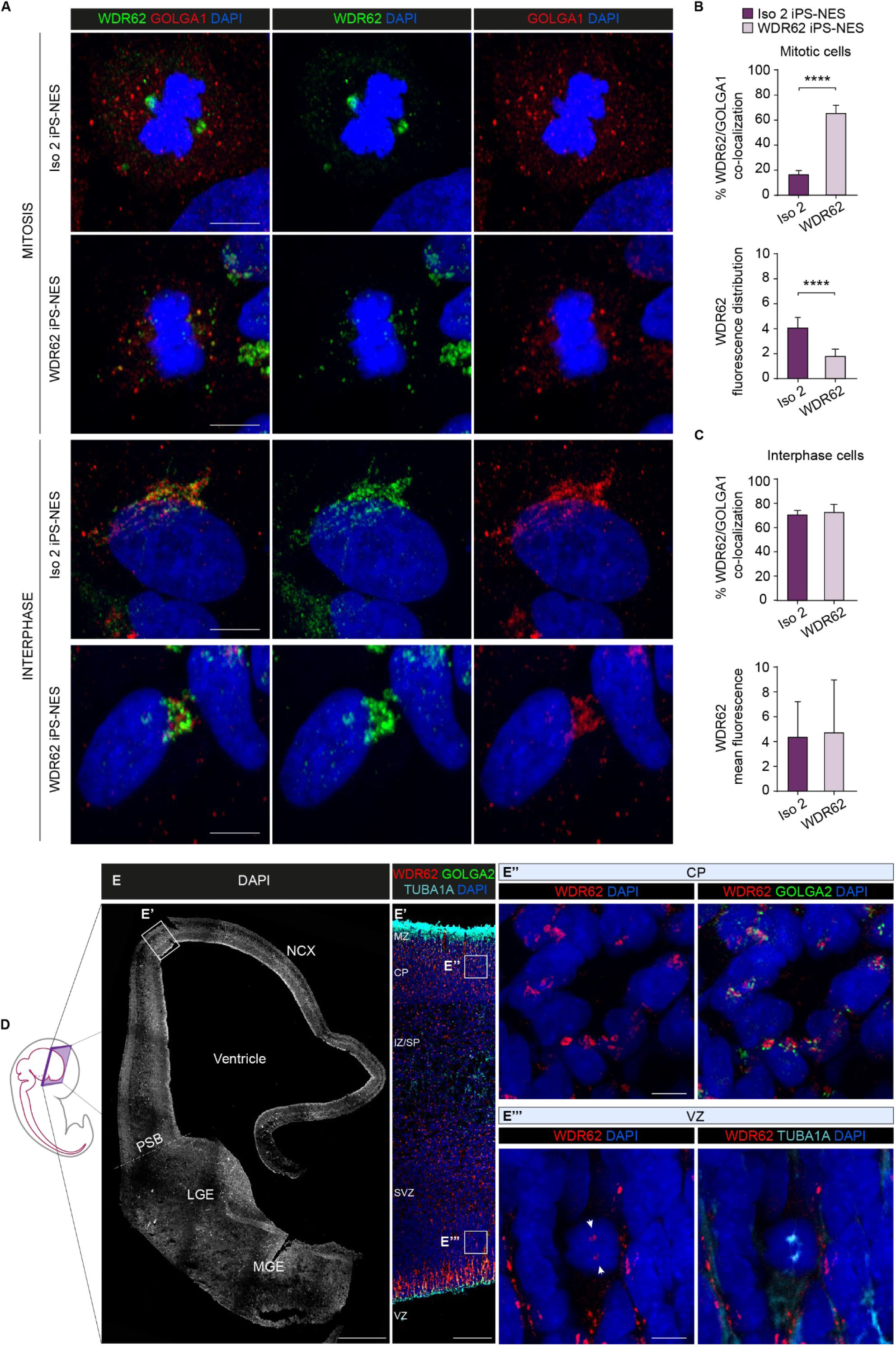
WDR62 displays cell cycle-dependent localization in Golgi/centrosome/spindle poles in iPS-NES cells and human fetal telencephalon. A) Immunofluorescence analysis of WDR62, GOLGA1, and TUBA1A in iPS-NES cells during mitosis (metaphase) and interphase. B-C) Analysis of WDR62 localization during cell cycle progression. In Iso 2 iPS-NES cells, WDR62 is localized to the spindle poles during mitosis, and co-localized with GOLGA1 in interphase. In WDR62 iPS-NES cells, the WDR62 signal remains diffuse and co-localizes with GOLGA1 during mitosis. The WDR62 subcellular localization is comparable in Iso 2 and WDR62 iPS-NES cells during interphase. In B) total cells N = 33, p-value < 0.0001, Kolmogorov-Smirnov test, top histogram; total cells N = 21, p-value < 0.0001, unpaired Student’s t-test, bottom histogram; fluorescence distribution, a.u.; in (c), total cells N = 60, p-value > 0.05, unpaired Student’s t-test, top histogram; total cells N = 164, p-value > 0.05, unpaired Student’s t-test, bottom histogram. D) Schematic illustration of a human fetal specimen showing the developing nervous system. E) DAPI staining of a coronal hemisection of human telencephalon at 9 post-conceptional weeks (pcw). E’) Magnification of the boxed area in E) showing WDR62/GOLGA2/TUBA1A expression in the developing NCX. E’’, E”’) Magnified views of the boxed areas in E’ showing WDR62/GOLGA2 expression in CP (E”) and WDR62/TUBA1A expression in VZ (E”’). Data are shown as mean ± SD. Scale bar = 10 μm in A, 500 μm in E, 100 μm in E’ and 5 μm in E’’’. NCX (Neocortex); PSB (Pallium-Subpallium Boundary); LGE (Lateral Ganglionic Eminence); MGE (Medial Ganglionic Eminence); MZ (Marginal Zone); CP (Cortical Plate); IZ/SP (Intermediate Zone/Subplate); SVZ (Sub-Ventricular Zone); VZ (Ventricular Zone).

We took several approaches to exclude that the WDR62 localization to the Golgi we observed could be due to lack of antibody specificity. First, we immunostained CTRL iPS-NES following siRNA-mediated knockdown of WDR62. No signal was detected in WDR62-silenced cells (Figure 3-figure supplement 3A-D). Next, we co-transfected CTRL iPS-NES cells with wild type or mutant hWDR62-FLAG and GALT1-mWasabi (which localizes to the Golgi (Cole et al., 1996)) (Figure 3-figure supplement 4A), and analyzed transfected cells with confocal imaging. We observed co-localization of FLAG (corresponding to wild type or mutant WDR62) and mWasabi signals, similar to what we observed for the endogenous WDR62 and Golgi signals (Figure 3-figure supplement 4B).

To strengthen these observations, we investigated WDR62 localization in sections of human fetal telencephalon from a presumed healthy subject at 9 post-conceptional weeks (pcw), when WDR62 is broadly expressed in neural progenitor populations (Bilgüvar et al., 2010; Nicholas et al., 2010). Strikingly, in the cortical plate, where committed neuroblasts are populating the incipient cortical layers, the WDR62 signal was polarized and perinuclear, overlapping with the Golgi marker GOLGA2 (also known as GM130), thus retracing the pattern observed in iPS-NES cells (Figure 3E’, 3E’’). In the cortical VZ, where radial glial progenitors reside and most mitotic events occur, WDR62 was visible at TUBA1A-positive spindle poles in mitotic cells (Figure 3E’, 3E’’’); in contrast, the WDR62 signal appeared elongated and scattered in interphase cells, reminiscent of the distribution of the Golgi apparatus along radial processes (Taverna et al., 2016) (Figure 3-figure supplement 5A-C). Finally, in human fetal incipient dorsal and ventral telencephalic areas at 11 pcw, WDR62 and GOLGA2 signals were polarized and perinuclear in the cortical plate and the mantle zone of the developing striatum, but scattered along NPC processes at the germinative zones (Castiglioni et al., 2019; Onorati et al., 2014) (Figure 3-figure supplement 5D-E).

Thus, these data suggest a dynamic association between WDR62 and the Golgi apparatus in human NPCs in vitro, and in the developing human telencephalon.

### WDR62 shuttling to spindle poles depends on microtubules

Next, we sought to investigate the mechanism underlying WDR62 shuttling from the Golgi apparatus to the spindle poles at the interphase/mitosis transition. As previously reported (Wang et al., 2010; Welburn & Cheeseman, 2012), several MCPH-related proteins translocate from various cellular compartments to the spindle poles or centrosomes along microtubular rails. Among them, CDK5RAP2, CEP63, together with WDR62 and ASPM, are known to belong to the same molecular assembly localizing hierarchically at the centrosome during interphase (Kodani et al., 2015).

First, we examined the consequences of microtubule depolymerization on WDR62 localization dynamics. Following nocodazole treatment and fixation, WDR62 appeared diffuse around the nucleus in mitotic Iso iPS-NES cells and was absent from the spindle poles or centrosome/pericentrosomal area, stained with CETN2 (Degl’Innocenti et al., 2022) or PCTN (Figure 4A); the same pattern was also observed in WDR62 iPS-NES cells (Figure 3B). We analyzed WDR62 signal intensity and distribution around chromatin and mean fluorescence values (Figure 4B) before and after nocodazole treatment, confirming that the observed pattern was comparable to mutant WDR62 distribution in untreated WDR62 iPS-NES cells (Figure 3B). Moreover, the co-localization percentage of WDR62 to the Golgi apparatus was similar in treated Iso and WDR62 iPS- NES cells (Figures 3B and 4B). Together, these observations strongly suggest that microtubules are fundamental for Golgi-to-spindle pole translocation of WDR62 during mitosis, and that the C-terminal part of the protein may have an important role in its interaction with microtubules.

**Figure 4:**
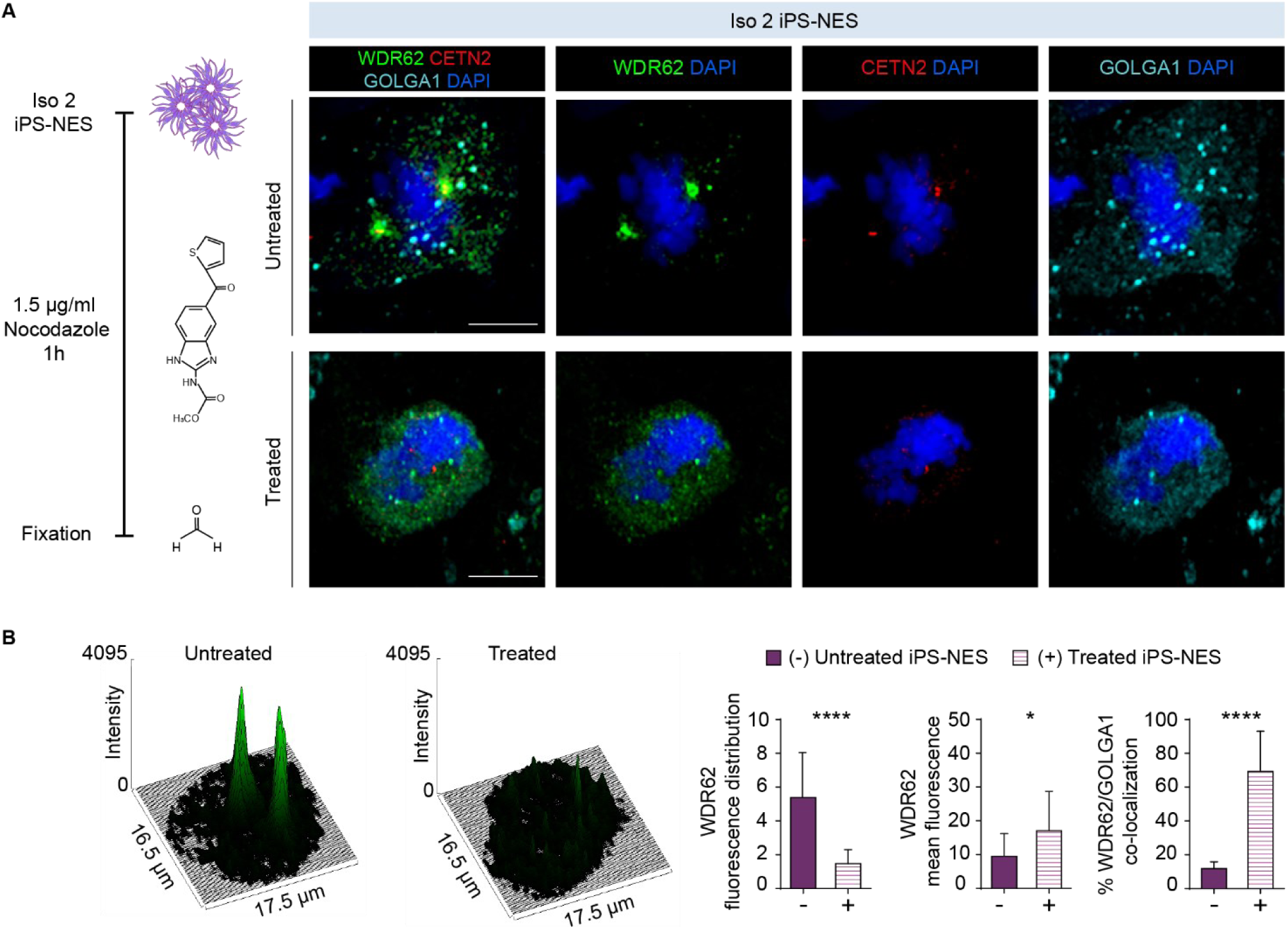
Microtubule-dependent shuttling of WDR62 to spindle poles. A) Immunofluorescence analysis of WDR62, GOLGA1, and CETN2 in Iso 2 iPS-NES cells after microtubule depolymerization (1.5 μg/ml nocodazole for 1 h). B) Analysis of WDR62 fluorescence reveals a similar signal distribution, mean fluorescence, and co-localization with GOLGA1 in nocodazole-treated Iso and WDR62 iPS-NES cells (see Figure 3A, B) during mitosis. Total cells N = 42, p-value < 0.0001, Kolmogorov-Smirnov test, left histogram; total cells N = 42, p-value < 0.05, Kolmogorov-Smirnov test, center histogram; total cells N = 32, p-value < 0.0001, unpaired Student’s t-test, right histogram. Data are shown as mean ± SD. Scale bar = 10 μm.

Then, we asked whether the centrosomal localization of CDK5RAP2 was preserved in WDR62 NES cells. We found that, both during mitosis and interphase, CDK5RAP2 was correctly localized, as we reported previously for skin fibroblasts (Sgourdou et al., 2017) (Figure 4-figure supplement 1A). Then, we investigated CDK5RAP2 localization in Iso iPS-NES cells after nocodazole treatment. We found that, already during G_1_ and before centriole duplication, CDK5RAP2 was localized to the centrosome both in untreated and nocodazole-treated cells (Figure 4-figure supplement 1B), whereas WDR62 remained localized to the Golgi (Figure 3B). After centriole duplication at early S-phase (Holland et al., 2010), CDK5RAP2 was detected at the centrosomes despite nocodazole treatment (O’Neill et al., 2022), even before centrioles started to migrate to participate in spindle pole formation (Figure 4-figure supplement 1B). During prometaphase, in untreated Iso iPS-NES cells, both CDK5RAP2 and WDR62 co-localized to the centrosome and properly positioned spindle poles, respectively (Figure 4-figure supplement 1B). In nocodazole-treated cells, however, the two proteins showed divergent behavior: CDK5RAP2 remained at the centrosome (Figure 4-figure supplement 1C), whereas WDR62 appeared dispersed (Figure 4-figure supplement 1D). WDR62 and CDK5RAP2 have been reported to physically interact (Kodani et al., 2015). We overexpressed CDK5RAP2 and wild type or mutant WDR62 in HEK293T cells and then performed co-immunoprecipitation (co-IP). We found that the mutation did not prevent WDR62 interaction with CDK5RAP2 (Figure 4-figure supplement 1E and Figure 4-figure supplement 1-source data 1). These findings support the hypothesis that the mutation does not impinge physical interaction with CDK5RAP2, but prevents WDR62 shuttling to the spindle poles. Moreover, we tested by co-IP, in the same overexpression system, the effect of the mutation on WDR62 binding with other known interactors AURKA and TPX2 (Lim et al., 2015, 2016) (Figure 4-figure supplement 1F, G and Figure 4-figure supplement 1-source data 2 and 3). Our results indicate that both interactions are preserved in the presence of mutant WDR62.

### The D955AfsX112 mutation impairs neurogenic timing

Considering its impact on self-renewing NES cells, we next investigated potential effects of the WDR62 mutation on iPSC differentiation into mature cortical neurons and glia. We employed a directed cerebro-cortical differentiation protocol (Figure 5A): after the initial neuroinduction phase, we obtained, at 16 days in vitro (DIV16), neuroectodermal cells, which were subsequently driven toward terminal differentiation through addition of supplements and neurotrophins (i.e., BDNF) (Chambers et al., 2009; Shi et al., 2012). Quantification of MKI67^+^ and PH3^+^ mitotic cells, as well as EdU-incorporating S-phase cells, revealed an increase in mitotic index (PH3^+^) in progeny of WDR62 versus Iso iPSCs at DIV16 (Figure 5B), but no differences at DIV40 (Figure 5C). Neuronal maturation, reflected by the percentage of MAP2^+^ cells, was similar in WDR62 and Iso iPSC-DIV40 (Figure 5C), suggesting that general neurogenic properties were not altered. Next, we investigated layer-specific fate of early-born neurons in DIV40 cultures and observed a significant enrichment in neurons expressing TBR1 and BCL11B (also known as CTIP2), markers of layer 5 and 6, respectively, in WDR62 compared to the Iso iPSC progeny (Figure 5C). Taken together, these results indicate that the mutation results in neuronal fate commitment imbalance, rather than a generalized maturation impairment.

**Figure 5:**
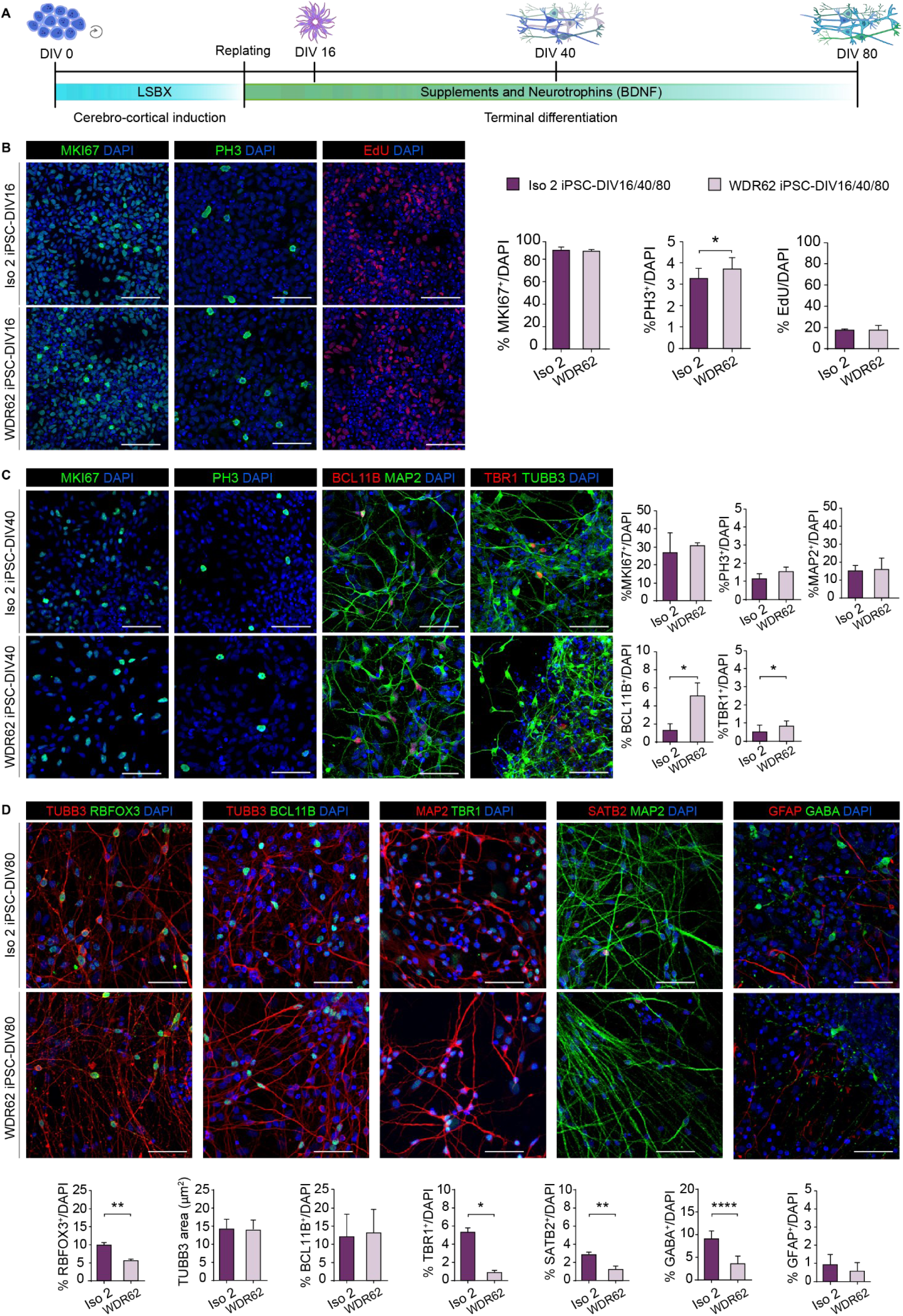
The D955AfsX112 mutation leads to lineage specification defects in iPSC-derived cerebro-cortical populations. A) Schematic illustration of the cerebro-cortical differentiation protocol. B) Immunofluorescence analysis for MKI67, PH3, and EdU in Iso and WDR62 iPSC neural progeny at DIV16. Quantitative analysis (right) shows no differences in the number of proliferating (MKI67^+^) or S-phase (EdU^+^) cells, but a significative increase in mitotic index (PH3^+^) in WDR62 iPSC neural progeny. Total cells N = 5222, p-value > 0.05, unpaired Student’s t-test for MKI67; total cells N = 58313, p-value < 0.05, unpaired Student’s t-test for PH3; total cells N = 3395, p-value > 0.05, unpaired Student’s t-test for EdU>50%. C) Immunofluorescence analysis for MKI67, PH3, BCL11B, MAP2, and TBR1 in Iso 2 and WDR62 iPSCs at DIV40 (early-born neurons; left). Quantitative analysis (right) shows no significant differences in the number of proliferating cells (MKI67^+^) or mitotic index (PH3^+^) in Iso 2 and WDR62 iPSC-progeny. BCL11B^+^ and TBR1^+^ neurons are significantly enriched in WDR62 compared to Iso 2 iPSC-DIV40 cultures. Quantification of MAP2^+^ cells did not show differences. Total cells N = 18795, p-value > 0.05, unpaired Student’s t-test, post hoc Welch’s correction for MKI67; total cells N = 15423, p-value > 0.05, unpaired Student’s t-test for PH3; total cells N = 40866, p-value < 0.05, unpaired Student’s t-test for BCL11B; total cells N = 38412, p-value < 0.05, unpaired Student’s t-test for TBR1; total cells N = 39595, p-value > 0.05, unpaired Student’s t-test for MAP2. D) Representative confocal images for RBFOX3, TUBB3, BCL11B, TBR1, SATB2, GABA, and GFAP staining in Iso 2 and WDR62 iPSCs at DIV80 (top). Quantification (bottom) shows a decrease in RBFOX3^+^, TBR1^+^ and SATB2^+^ cells in WDR62 iPSC derived neurons, and a decrease in GABA^+^ but not in GFAP^+^ cells or neurite area (TUBB3^+^). Total cells N = 35207, p-value < 0.01, unpaired Student’s t-test for RBFOX3; total fields measured, N = 6, p-value > 0.05, unpaired Student’s t-test for TUBB3; total cells N = 29448, p-value > 0.05, unpaired Student’s t-test for BCL11B; total cells N = 23396, p-value < 0.05, unpaired Student’s t-test for TBR1; total cells N = 21843, p-value < 0.01, unpaired Student’s t-test for SATB2; total cells N = 20375, p-value < 0.0001, unpaired Student’s t-test for GABA; total cells N = 20370, p-value > 0.05, unpaired Student’s t-test for GFAP. Data are shown as mean ± SD. Scale bars: 10 μm.

We then investigated neuronal lineage composition at DIV80. We analyzed the general maturation state by staining with the pan-neuronal marker RBFOX3 (also known as NeuN) and TUBB3, to estimate total neurite-occupied area, an indicator of neuronal differentiation (Figure 5D). Compared with Iso counterparts, WDR62 iPSC progeny contained fewer RBFOX3^+^ neurons, even in the absence of noticeable differences in the neurite-occupied area (Figure 5D). We also examined overall layer identity at DIV80 by quantifying neurons expressing TBR1 and BCL11B. Although the percentage of BCL11B^+^ cells was similar in WDR62 and Iso iPSC progeny (thus compensating the early imbalance), we observed a significant reduction in TBR1^+^ cells in the former, reversing the premature differentiation observed at DIV40 (Figure 5D). We further noticed a decrease in SATB2-expressing layer 2/3 neurons in WDR62-iPSC progeny (Figure 5D). Lastly, we observed a reduction in GABAergic interneurons, but a normal proportion of GFAP-expressing astrocytes in WDR62-iPSC derivatives (Figure 5D). Together, these results strongly suggest neural lineage alterations associated with the WDR62 mutation and a subtle maturation delay during differentiation, potentially recapitulating the neocortical layering abnormalities previously observed in MCPH patients and mouse models (Sgourdou et al., 2017).

## Discussion

WDR62 is a spindle pole associated protein with key functions in NPC mitotic progression during cortical development. Recessive mutations in WDR62 account for the second most common cause of MCPH, a genetic neurodevelopmental disorder characterized by a spectrum of severe brain malformations and intellectual disability (Bilgüvar et al., 2010; Nicholas et al., 2010; Yu et al., 2010).

In this study, we investigated the impact of a C-terminal truncating mutation (D955AfsX112) in WDR62 (Sgourdou et al., 2017) on human cortical development employing patient iPSCs, which we used to derive NES cells and, through directed differentiation, mature cerebro-cortical neurons, and glia (Abdullah et al., 2017; Baggiani et al., 2020; Dell’ Amico et al., 2021). To date, human iPSC-derived NES cells and neurons have been employed in mechanistic studies of several neurodevelopmental disorders, including the CEDNIK syndrome, characterized by cerebral dysgenesis (Morelli et al., 2021), Machado–Joseph disease (spinocerebellar ataxia 3) (Koch et al., 2011), and Zika virus-induced microcephaly (Lottini et al., 2022; Onorati et al., 2016) . NES cells have also been explored as means of cell therapy approaches after spinal cord injury (Dell’Anno et al., 2018).

Here, through analyses of iPSC-derived NES cells and mature neurons harboring the D955AfsX112 mutation and isogenic-corrected counterparts, we found that: i) mutant WDR62 fails to localize to the spindle poles during mitosis; ii) patient-derived iPS-NES cells exhibit shorter primary cilia and significantly smaller spindle angles, defects that likely contribute to the previously observed impairment in cell cycle progression; iii) the mutation leads to lineage specification defects in iPSC-derived cerebro-cortical neurons; iv) during the interphase-to-mitosis transition, WDR62 translocates from the Golgi apparatus to the spindle poles in a microtubule-dependent manner; and v) pharmacological disruption of microtubules mimics the effects of the genetic mutation and prevents WDR62 shuttling from the Golgi to the spindle poles. WDR62 contains a WD40-repeat region at its N-terminus and several regulatory domains at the C-terminal half and has cell cycle dependent functions. Even though AURKA-mediated phosphorylation of its N-terminus is required for WDR62 accumulation to the spindle poles, spindle pole localization is disrupted both by N-(Lim et al., 2015) and C-terminal (Farag et al., 2013; Nicholas et al., 2010) mutations. The D955AfsX112 truncating mutation (Sgourdou et al., 2017) investigated in this study is expected to disrupt the AURKA binding domain (aa 621-1138) (Chen et al., 2014), as well as the JNK1 phosphorylation site (T1053) previously reported to diminish WDR62 association with the microtubules (Lim et al., 2015). Here we show that, in mitotic WDR62 iPSCs as well as in WDR62 iPS-NES cells, WDR62 fails to localize to the spindle poles, in line with other reports (Farag et al., 2013; Nicholas et al., 2010). The interaction with AURKA, however, appears preserved, even if weaker, at least in a heterologous system upon overexpression. Importantly, restoration of the correct WDR62 sequence through genome editing reverted the mutant phenotype, allowing relocation of WDR62 to the spindle poles during mitosis, and supporting the causal association between the identified mutation and the phenotype.

Since nocodazole-mediated microtubular disruption mimics in Iso iPS-NES cells the pattern observed for mutant WDR62, we are prompted to assume that the C-terminus likely regulates microtubules interaction, which is required for WDR62 translocation to the spindle poles. Indeed, although microtubule binding is thought to be mediated by the N-terminus (Lim et al., 2015), our work suggests that the C-terminus is also needed, even if the mechanism remains unknown. This is also in agreement with observations by Lim et al., 2015 (Lim et al., 2015) that the C-terminus MCPH-associated mutations fail to localize WDR62 to spindle microtubules.

Here, we also provide evidence for WDR62 localization at the Golgi apparatus during interphase in human WDR62 as well as Iso iPS-NES cells and in human fetal telencephalon, reminiscent of previous observations in HeLa cells (Nicholas et al., 2010; Yu et al., 2010). Notably, several syndromes caused by genetic defects affecting Golgi-related proteins present with microcephaly, corpus callosum agenesis or thinning, and neurological defects (Banne et al., 2013; Dimitrov et al., 2009). Moreover, the disruption – especially during neurogenic processes – of Golgi-associated proteins which are also involved in cell cycle regulation or cellular polarity/microtubule organization lead to disorders underlain by lack of neurons/neuronal migration defects (e.g., MCPH3 (Wang et al., 2010) and MCPH17 (Camera et al., 2008; H. Li et al., 2016)).

Even though the mechanism(s) leading to loss of spindle pole localization are yet to be precisely determined, our findings in WDR62 iPS-NES cells indicate a general cell cycle regulation impairment and cell cycle re-entry failure in neural progenitors. This altered behavior becomes more evident after inducing cell cycle arrest. Taken together with similar observations in patient skin fibroblasts (Sgourdou et al., 2017), our findings provide support for the role of WDR62 in cell cycle regulation and, by extension, neocorticogenesis. Our findings of smaller spindle angle in WDR62 iPS-NES cells may in turn reflect defective control and timing of cell fate determination (Fish et al., 2006; Lesage et al., 2010; Lizarraga et al., 2010). In addition, we report that WDR62 iPS-NES cells harbor shorter primary cilia, which are known to play important roles in cortical development (Liu et al., 2021). Indeed, shorter cilia could explain the disrupted mitotic progression we observed in WDR62 iPS-NES cells (Kasahara & Inagaki, 2021). Zhang and colleagues (Zhang et al., 2019) previously reported interactions of the N-terminus of WDR62 with the CEP170-KIF2A pathway in promoting cilium disassembly in neural progenitors. The longer primary cilia these authors observed in WDR62 KO cerebral organoids (Zhang et al., 2019) could be due to the complete deletion of WDR62, which has pleiotropic actions. We speculate that the shorter primary cilia we observe in WDR62 iPS-NES cells might reflect disruption in the assembly, rather than the disassembly of the primary cilium, considering that the mutation under study results in C-terminal truncation (Shohayeb et al., 2020).

It is not surprising that cell cycle impairment in neural progenitors constitutes the substrate for altered neurogenic processes, ultimately resulting in MCPH (Jayaraman et al., 2018). Indeed, during directed cerebro-cortical differentiation, we observed increased mitotic index in WDR62 iPSC neural cultures at DIV16. Moreover, we observed an enrichment in BCL11B^+^ and TBR1^+^ cells in WDR62 iPSC-DIV40 neurons, indicating altered neurogenic trajectories. Surprisingly, this did not appear to be coupled with premature neuronal maturation (measured by MAP2 positivity), suggesting incorrect cell fate acquisition/neurogenic timing rather than generalized premature differentiation. Of note, late progenitors were reported to be primarily affected in a mouse model homozygous for a trapped Wdr62 allele leading to the expression of a fusion WDR62 (a.a. 1-870)-beta-galactosidase protein (Sgourdou et al., 2017).

At late stages (DIV80), the WDR62 iPSC-neuronal progeny consisted of fewer overall mature (RBFOX3^+^) neurons, as well as fewer deep (TBR1^+^) and upper (SATB2^+^) layer neurons; however, the earlier imbalance in the number of BCL11B^+^ and TBR1^+^ neurons appeared to have resolved. Further hints of neurogenesis impairment can be found in GABAergic neuron population, which is reduced in WDR62 iPSC-DIV80. However, neither the area of neurite occupancy, which includes length and neurite number, nor gliogenesis, appear to be affected by the WDR62 mutation. In contrast, the composition of Iso iPSC neural progeny was as expected and in agreement with previous reports (Shi et al., 2012).

Recently, Vargas-Hurtado et al., 2019 (Vargas-Hurtado et al., 2019) found that at early neurogenic stages, mitotic spindles display more astral microtubules and fewer spindle microtubules, whereas the opposite was observed at late neurogenic stages. Additionally, it has also been speculated that during neocortical development the centrosomal microtubule-organizing center (MTOC) undergoes different activation states during the transition from radial glial cells to migrating neuroblasts (Camargo Ortega et al., 2019; Camargo Ortega & Götz, 2022). Since WDR62 localizes to the Golgi apparatus, which is interconnected with centrosomal MTOC itself (Bornens, 2021), and physically interacts with CDK5RAP2/CEP170/CEP63 (Kodani et al., 2015), localized at the centrosomes of neural stem cells (O’Neill et al., 2022) (Huang et al., 2021) it is indeed reasonable to speculate that during the transition from early to late neurogenic stages, WDR62 is involved in the switch in spindle morphology and in the regulation of neural stem cell differentiation, through interactions with centrosomal MTOC proteins.

Thus, loss of function of WDR62 may cause altered neurogenic trajectories at the different stages, eventually altering the final neuronal output observed in the MCPH associated with WDR62 mutations.

Together, these findings indicate that the D955AfsX112 mutation in WDR62 results in incorrect neuronal cell fate commitment secondary to neural progenitor defects, in line with other reports (Sgourdou et al., 2017; Zhang et al., 2019). Our study also provides evidence that WDR62 is localized to the Golgi apparatus during interphase in human iPS-NES cells in vitro as well as in the developing human forebrain, reminiscent of previous reports in HeLa cells (Nicholas et al., 2010; Yu et al., 2010). A physical interaction of WDR62 with the Golgi protein GOLGA6L2 has also been reported in Bioplex, a database of human protein-protein interactions generated by affinity purification mass spectroscopy (Huttlin et al., 2015, 2017, 2021).

Our observations suggest a dynamic pattern of WDR62 shuttling from the Golgi to the spindle poles which is microtubule-dependent, and impaired upon microtubule de-stabilization by nocodazole. Our findings also suggest that the mutation impairs normal shuttling from the Golgi apparatus, thus explaining the lack of WDR62 localization at the spindle poles during mitosis. Of note, the D955AfsX112 mutation does not appear to impair the WDR62 interactions with spindle pole binding partners, supporting the hypothesis that shuttling from the Golgi apparatus is microtubule dependent.

In conclusion, the study of a C-terminal truncating mutation through iPSC-derived models allowed an in-depth and broader appreciation of the molecular mechanisms underlying WDR62-related microcephaly and, more generally, of MCPH etiology.

## Materials and Methods

### Ethical statement

All cell work was performed according to NIH guide-lines for the acquisition and distribution of human tissue for biomedical research purposes and with approval by the Human Investigation Committee and Institutional Ethics Committee of each institute from which the samples were obtained (University of Pisa Review No. 29/2020 and Yale No. 9406007680). Appropriate informed consent was obtained, and all available non-identifying information was recorded for each specimen. The tissue was handled in accordance with the ethical guidelines and regulations for the research use of human brain tissue set forth by the NIH and the WMA Declaration of Helsinki. The human fetal brain sections used in this study were previously obtained by Bocchi et al., 2021 (Bocchi et al., 2021) and derived from post-mortem specimens as approved by University of Cambridge and Addenbrooke’s Hospital in Cambridge (protocol 96/85).

### Description of the mutation

The mutation under study was identified in the index case NG1406-1 and described in Sgourdou et al., 2017 (Sgourdou et al., 2017). Briefly, analysis of whole exome sequencing data detected a novel homozygous frameshift mutation, D955AfsX112, caused by a 4 bp deletion in exon 23 of WDR62. This mutation leads to the generation of a premature stop codon at position 1067 producing a C-terminally truncated peptide (Sgourdou et al., 2017).

### iPSC culture and maintenance

Reprogramming of patient-derived fibroblasts (NG1406-1) was performed via episomal reprogramming vectors - pCXLEhOCT3/4-shp53-F, pCXLE-hSK, pCXLE-hUL (Sgourdou et al., 2017). Human iPSCs were cultured as previously described (Sousa et al., 2017). Briefly, cells were seeded on Matrigel-coated culture plates (Cat: #356234, Corning, 1:60) and maintained in StemFlex Basal medium (Thermo Fisher Scientific; # A3349201) or Essential 8^TM^ Medium (Thermo Fisher Scientific, # A2858501). Cells were typically passaged with EDTA (0.5 mM) at room temperature (RT). After 3-5 min of incubation, the EDTA solution was removed and cells were gently detached from the dish with a small volume of medium, generating clumps of 6-8 cells. In standard conditions (37°C, 5% CO_2_), iPSC colonies typically grow within 4-5 days.

### CRISPR/Cas9 genome editing: isogenic line generation

CRISPR/Cas9 technology was used to generate isogenic lines (Skarnes et al., 2019): 1.5x10^6^ WDR62 iPSCs (clone 1b2-1) were nucleofected with pre-mixed 8 µg of synthetic chemically-modified sgRNA_1 (Synthego), 200 pmol/µl of single strand donor oligonucleotide (ssODN; IDT) together with 20 µg of HiFi Cas9 Nuclease V3 (Cat: #1081060, IDT) using Amaxa™ 4D-Nucleofector™ (Cat: #AAF-1002B, Lonza) transfection system with Primary Cell Optimization 4D-Nucleofector X Kit (Cat: #V4XP-9096, Lonza) in P2 buffer – program DS-150.

After nucleofection, cells were counted and assessed for viability with trypan blue (cell death around 18%), seeded onto Matrigel-coated culture plates supplemented with 10 µM Y-27632 (Cat: #72308, StemCell Technologies) and incubated at 32°C (5% CO_2_) for 2 days. This passage was instrumental to increase the rate of homologous recombination (Guo et al., 2018). The day after nucleofection, cell medium was replaced to remove Y-27632. Two days later, cells were detached and plated at low density in 60 cm^2^ Petri-dishes pre-coated with Matrigel and cultured for 15 days, until cell-colonies were ready to be picked. The three clonal iPSC lines Iso 1, Iso 2, and Iso 3 were selected and fully characterized. No interclonal variability was observed in the different assays. The specific Iso iPSC line used for each assay is indicated the figures.

To assess WDR62 sequence restoration, single clonal populations were first screened through melting curve analysis (Erali et al., 2008) performed on a CFX96-BioRad cycler. WDR62_F1 and WDR62_R1 primers were used in combination and samples were prepared using the SensiFast^TM^ HRM kit for RT-qPCR - melt curve reaction as follows. Amplification phase: Initial denaturation: 95°C 5s; Denaturation: 95°C 10s; Annealing: 60°C 20s; Extension: 72°C 20s; Cycles number: 40. Melting phase: 95°C 30s. Melt curve 65°C to 95°C, increment 0.1°C – 5s.

The relevant portion of genomic DNA from selected clones was amplified by PCR using WDR62_GF1 and WDR62_GR1 primers, with the following conditions: Initial denaturation: 95°C 2 min; Denaturation: 98°C 20s; Annealing: 55°C 30s; Extension: 72°C 1 min; Final extension: 72°C; 5 min. Cycle number: 30. Finally, the genotype was confirmed by Sanger sequencing. Sequences for genome editing (5’-3’): sgRNA_1: GAAGTGACAGTCACAGGGAC; ssODN: GCCAGTGAGCTCATCCTCTACTCTCTGGAGGCAGAAGTGACAGTCACAGGGACAG<u>ACAG</u>GTG GGTGTCCTTTCCACAAGGGAGCCTTAGTTGGAGGAACCCCCAGCTG Primer sequences (5’-3’): Melting analysis: F1: CTCTGGAGGCAGAAGTGACAG R1: CTTGGTGGAAAGGACACCCAC. PCR: GF1: ACTGGGTTTCCTATTCTTGAACTTG GR1: AGGACTTCAGCTGGAGACTCAAC. Sequencing: SF1: TGTGCTGTCTTCCCCATAGTC; SR1: CCCATCCAGGCCTCAACTGTC

### Generation of cerebro-neocortical neurons

Mature cortical neurons were generated from iPSCs through an optimized cerebro-cortical differentiation protocol, based on a dual SMAD inhibition strategy (Chambers et al., 2009; Shi et al., 2012). As previously described (Sousa et al., 2017), iPSCs were dissociated into a single-cell suspension with Accutase (Corning, #25-058-CI) pre-warmed at 37°C, and plated onto Matrigel-coated 6 well-plates at high density (0.7 - 2 x 10^5^ cells/cm^2^) in StemFlex basal media supplemented with Y-27632 (10µM). When they reached a confluent state, maintenance medium was replaced with neural induction medium [1:1 DMEM/F-12 (Thermo Fisher Scientific, #31330095) and Neurobasal^TM^ medium (Thermo Fisher Scientific, #21103049) supplemented with N-2 (1:100, Gibco, #17502-048), B-27 (1:50, Thermo Fisher Scientific, #17504-044), 20 µg/ml insulin (Sigma-Aldrich, #I9278-5ML), L-glutamine (1:100, Thermo Fisher Scientific, #25030-081), MEM non-essential amino acids (1:100, Gibco, #11140-050) and 2-mercaptoethanol (1:1000, Thermo Fisher Scientific, #31350010). For cerebro-cortical induction (LSBX), 100 nM of LDN-193189 (STEMCELL Technologies, #72144), 10 µM of SB-431542 (Merck, #616464-5MG) and 2 µM of XAV939 (STEMCELL Technologies, #72674) were added to induce cerebro-cortical differentiation. Induction medium was replaced daily until day 11. For terminal differentiation, cells were dissociated at day 12 with Accutase and replated onto Poly-D-Lysine (Sigma, #P6407, 10 µg/ml)/Laminin (Invitrogen, #23017-015, 3 µg/ml) coated chamber slides at the density of 8 x 10^4^ cells in terminal differentiation medium containing Neurobasal^TM^ supplemented with N-2 (1:100), B-27 (1:50), L-glutamine (1:100), BDNF (30 ng/ml, R&D Systems, #248-DB025), and Y-27632 (10 μM) to increase cell viability. Culture media were partially replaced every 3-4 days until day 80.

### NES cell generation

To obtain NES cells, at the end of the neuroinduction phase (day 12), 2x10^6^ cells were replated onto POLFN-coated 6-well plates (0,01% of Poly-L-ornithine, Sigma, #P4957; 5 µg/ml Laminin; 1 µg/ml Fibronectin, Corning, #354008) in NES medium containing DMEM/F-12 supplemented with B-27 (1:1000), N-2 (1:100), 20 ng/ml of FGF-2 (Gibco, #13256 029), 20 ng/ml of EGF (Gibco, #PHG0311), glucose (1.6 g/L), 20 µg/ml insulin and 5 ng/ml of BDNF. NES cells were then cultured as previously reported (Dell’Anno et al., 2018); to preserve optimal growth properties, the medium was replaced every 2-3 days and cells passaged 1:2 or 1:3 weekly with 0.25% Trypsin/EDTA (Thermo Fisher Scientific, #25200-056) pre-warmed at 37°C.

### NES cell synchronization

For efficient synchronization, NES cells were seeded onto POLFN-coated coverslips in a 48-well plate at a density of 10^5^ cells/cm^2^. To induce prometaphase arrest (T1), cells were rinsed once with D-PBS (Sigma-Aldrich, # D8537) and cultured in NES medium containing 100 ng/ml nocodazole (Millipore Sigma, #M1404) for 16 h. Then, cells were fixed with 4% formaldehyde (FA) for 20 min at 25°C.

For cell cycle re-entry analysis (T2), after 16 h of treatment, nocodazole was removed and synchronized NES cells were rinsed once with D-PBS, released in NES medium for 1 h and then fixed in 4% FA for immunofluorescence analysis. For mitotic index and mitotic figure distribution analysis cells were classified following PH3 staining.

### Microtubule de-polymerization assay

To induce microtubule de-polymerization, NES cells were first plated onto POLFN-coated coverslips in a 48-well plate at the density of 10^5^ cells/cm^2^. The day after, 5 µM nocodazole was added to cell culture medium for 1 h; the cells were then rinsed once with D-PBS and fixed (4% FA for 20 min at 25°C) for immunofluorescence analysis.

### Mitotic spindle angle and inter-centrosomal distance estimation

Confocal images of NES cells immunostained with the pericentrosomal marker PCTN were acquired with a pre-determined Z-stack step-size of 300 nm and elaborated with ImageJ software (Schneider et al., 2012) (developed by National Institute of Health; https://imagej.nih.gov/ij). Mitotic spindle angle (θ) was calculated as follows: θ = arctan (h/D), with “h” as the distance between centrosomes on Y-axis and “D” as the distance between centrosomes on X-axis, manually traced (edge-to-edge distance of PCTN signal). Inter-centrosomal distance (ICD) was calculated as follows: ICD = D/cos (θ) (see Figure 2D for reference).

### Primary cilium length

Serial optical Z-stack with a step-size of 300 nm were used to create Z-projections (maximum intensity projection) of primary cilia for a minimum of 50 cells per genotype. Single cilium length was measured using “NeuronJ” plugin of Image J (Meijering et al., 2004). Briefly, after selecting the first pixel mid-point at the cilium peri-nuclear edge, short subsequent segments were traced manually following cilium shape until the last pixel mid-point. Cilia were measured regardless of cell cycle phase, excluding mitosis.

### Quantitative real-time PCR

RT-qPCR was performed to quantify WDR62 mRNA in Iso and WDR62 iPS-NES cells cultures. iPS-NES cells were plated in 6-well plates and lysates were collected in RNAprotect® Cell Reagent (QIAGEN, #76526) to stabilize the RNA. Subsequently, RNA purification was performed applying RNAeasy® Protect Cell Mini Kit (QIAGEN, #74134) and RNase-Free DNase set (QIAGEN, #79254). cDNA was synthetized using GoScriptTM Reverse Transcription System (Promega, #A5001) according to the manufacturer’s instructions. PCR was performed using QuantStudioTM 3 Real-Time PCR System (Applied BiosystemsTM, A28137) with SensiMixTMSYBR®No-ROX kit (Meridian BIOSCIENCE, #QT650-05). Thermal cycling conditions: denaturation at 95°C for 10 min and 40 cycles of 95°C for 15s and 60°C for 1 min. Data are expressed as fold change in WDR62 gene expression relative to GAPDH housekeeping gene, according to the 2^−ΔΔCT^ method. Three technical replicates were performed for each experiment. qPCR primers (5’-3’) qF_WDR62: GGAGGAAGAGTGTGAGCCAG; qR_WDR62: CTTGCCGTTGGTTAGCAGG.

### EdU labelling

The Click-iT EdU AlexaFluor 594 Imaging kit (Invitrogen, #C10339) was used to perform EdU labelling on iPSC cultures at DIV16, according to the manufacturer’s instructions. 3.5 h after EdU incubation, cells were fixed, imaged with confocal microscopy, and processed using ImageJ. For quantitative analysis, cells with signal intensity above 50% were considered.

### Immunostaining assay

After cell fixation (4% FA for 20 min at 25°C or methanol for 7 min at -20°C for the CDK5RAP2 antibody), 3 washes with D-PBSX (1% vol/vol TritonX-100 - Sigma #T9284-500ml - in D-PBS 1x) were performed. Then, samples were incubated at RT in permeabilization solution (5% vol/vol TritonX-100 in D-PBS Ca^2+^/Mg^2+^) for 10 min and then in blocking solution (5% FBS, 3% vol/vol TritonX-100 in PBS Ca^2+^/Mg^2+^) for at least 1 h (RT). Subsequently, samples were incubated in the antibody solution (3% FBS, 2% vol/vol TritonX-100 in D-PBS Ca^2+^/Mg^2+^) with primary antibodies overnight (ON) at 4°C. The next day, samples were washed 3 times with D-PBSX before incubation with the corresponding secondary antibodies (all diluted 1:500) and DAPI (Sigma, #32670-25mg) for 1 h (RT). Samples were mounted with Aqua-Poly/Mount (VWR, #87001-902) on microscope slides for confocal microscopy.

For immunohistochemistry, human fetal brain sections were first thawed at RT, washed once with D-PBS for 10 min and permeabilized with 0.5% Triton X-100 in D-PBS for 10 min. Sections were then washed in D-PBS and treated with Sodium Citrate based-R-buffer A (EMS, #62706-10) in 2100-Retriever (EMS, #62706) at 120°C for 20 min. After antigen retrieval, the sections were washed with D-PBS and blocked with 5% horse serum (Thermo Fisher, #26050070), 1% BSA in D-PBS with 0.3% Triton X-100 for 1 h (RT). Primary antibodies were diluted in blocking solution and incubated at 4 °C ON. The following day the sections were washed three times in D-PBSX (1% vol/vol TritonX-100 in D-PBS 1x). Secondary antibodies and DAPI were diluted in blocking solution for 1 h (RT). Then the sections were washed twice in D-PBSX and once in D-PBS and finally mounted with Aqua-Poly/Mount on microscope slides for confocal microscopy.

### Co-localization analysis

Following immunofluorescent staining, co-localization analysis for WDR62 and GOLGA1 (also known as Golgin97) signals was performed using the DiAna plugin (Gilles et al., 2017) (ImageJ) on 300 nm Z-stack confocal images. This plugin allows to calculate co-localization percentage between two selected objects measuring the distance (on 3 dimensions) of every single pixel unit – defined by size through threshold setting – constituting the object itself. To calculate WDR62-Golgi co-localization, Golgi objects identified by GOLGA1 signal (“object A”) were selected as Regions of Interest (ROI) and the same ROI has been identified on the WDR62 channel (“object B”). Then, for each A and B object pair, co-localization of A volume on B was calculated. Images were acquired with a Nikon A1 Confocal Microscope and NIS-Elements AR 4.20.03 64-bit software. Confocal images were then processed with ImageJ and Adobe Photoshop 2020.

### Cell culture, transfection, immunoprecipitation, and Western blotting

To generate the expression vectors used for co-IP and over-expression experiments, full-length cDNA fragments encoding Myc-hCDK5RAP2, Myc-hAURKA, Myc-hTPX2, or hWDR62-FLAG, were amplified by PCR and inserted into a CAG-promoter plasmid backbone by In-Fusion HD (Takara). A plasmid encoding hWDR62-FLAG was generated by amplifying hWDR62-FLAG by PCR; the fragment was excised using XhoI and NotI and ligated into a CAG-promoter plasmid. A plasmid encoding hWDR62D955A-FLAG was generated by overlap extension PCR cloning using two products (a.a. 1-955 of hWDR62D955A and 955-1067 of hWDR62D955A-FLAG), excised using XhoI and NotI, and ligated into a CAG-promoter plasmid. The plasmid expressing GALT1-mWasabi was generated by inserting the fragment corresponding to amino-acids 1-61 of hB4GALT1-mWasabi into a CAG-promoter plasmid.

HEK293T cells, maintained in DMEM/10% FBS, were transiently transfected using Lipofectamine 3000 (Invitrogen) according to the manufacturer’s instructions. Cell lysates were prepared 24 h after transfection using lysis buffer [Tris-HCl 50mM (pH7.4), 1mM EDTA,150 mM NaCl, 1% NP40, and protease inhibitors]. For immunoprecipitation, cell lysates were incubated with FLAG antibody for 1 h at 4°C, followed by incubation with Dynabeads Protein G (Invitrogen # 10003D) ON at 4°C. The immune precipitates were washed three times with cold TBST and analyzed by Western blotting.

Over-expression experiments were performed through forward transfection. CTRL iPS-NES cells, maintained in NES medium without antibiotics, were transiently transfected using Lipofectamine 2000 (Invitrogen) according to the manufacturer’s instructions. Cells (0.7 x 10^5^) were co-transfected with 0.5 µg of hWDR62-FLAG (or hWDR62*D955A-FLAG) and GALT1-mWasabi in 1:1 molar ratio. 48 h after transfection cells were fixed with 4% FA at 25°C for 20 min and imaged with confocal microscopy.

### Cell culture, transfection, and Western blotting

1x10^6^ iPS NES cells, maintained in NES medium without antibiotics, were transiently transfected using Lipofectamine 2000 as described above. Cells were lysed 48 h after transfection using lysis buffer [RIPA buffer (Sigma, #R0278), protease (Roche, #11836170001), and phosphatase (Roche, #04906837001) inhibitors]. After DC assay (Bio-Rad, # 5000111) for protein quantification, samples were analyzed by Western blotting (anti-WDR62 and anti-TUBA1A antibodies).

### Quantitative and statistical analysis

Nuclei and nuclear markers were quantified manually with the “Cell counter” plugin (ImageJ) or automatically through the “Analyze particles” function (ImageJ) defining a 15 µm^2^ particle-size threshold. Values identify mean and error bars represent standard deviation (S.D.) and a significance level of at least p < 0.05 (*p < 0.05; **p < 0.01; ***p < 0.001; ****p < 0.0001). All experiments were performed at least in triplicate. The size of the population (n) is reported for each experiment in the figure legend. Counts and analyses were performed blinded to conditions or genotypes. Graphs were generated with GraphPad Prism 7.00 software.

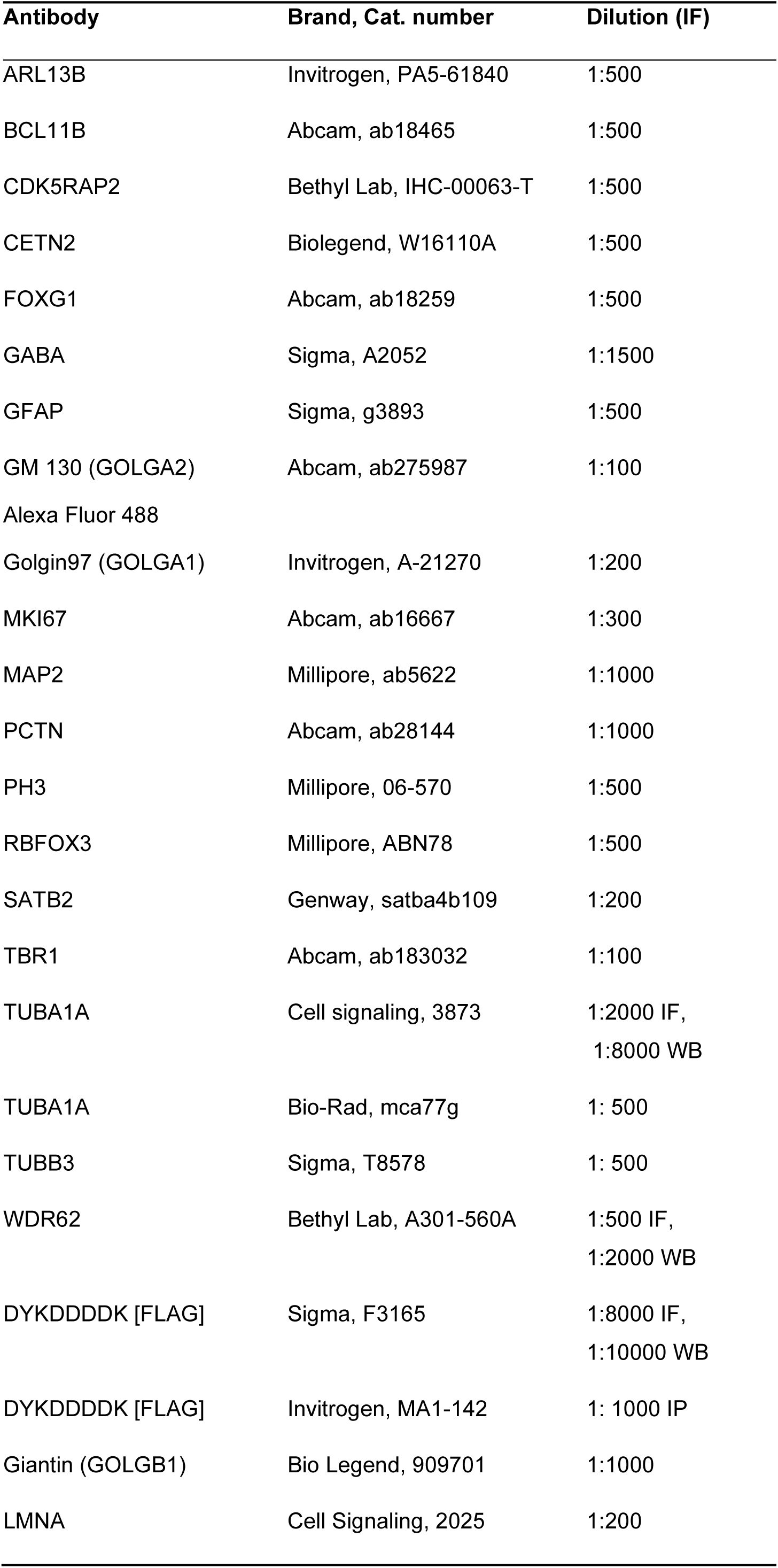
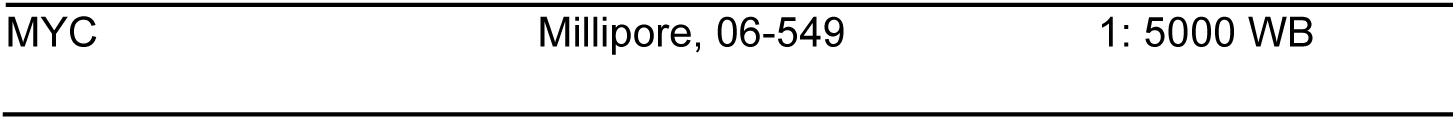

## Acknowledgements

We thank Catello Guida, Camilla Focacci, and Guglielma De Matienzo (University of Pisa, Department of Biology) for technical support and imaging analysis and Francesco Olimpico (Fondazione Pisana per la Scienza) for assistance in cell culture.

---

**Figure 1-figure supplement 1:**
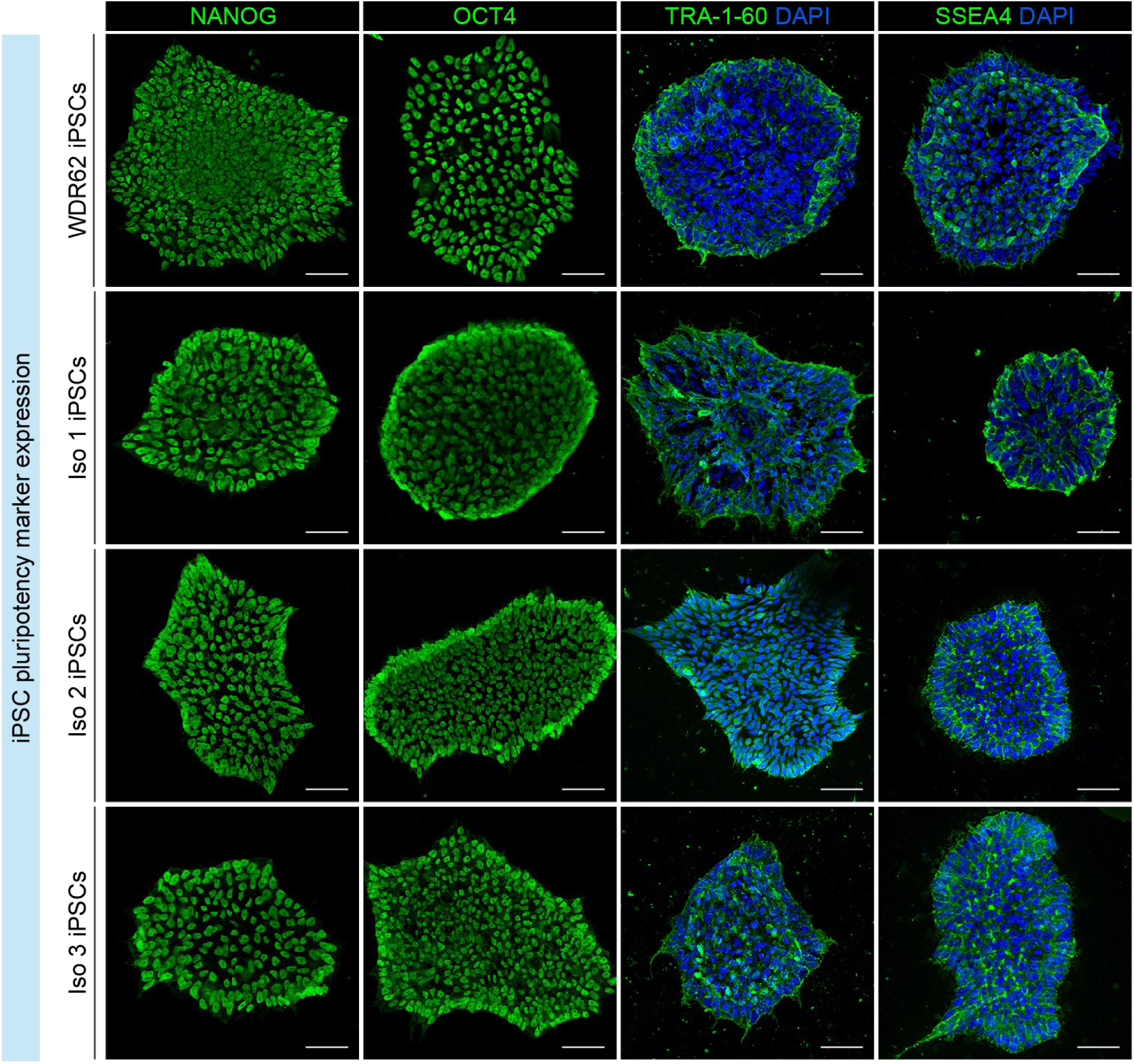
Analyses of pluripotency marker expression in mutant WDR62 and Iso iPSC lines. Immunofluorescence analysis for NANOG, POU5F1 (also known as OCT4), TRA-1-60, and SSEA4 in WDR62 and isogenic (Iso) iPSC lines. Scale bar = 50 μm.

**Figure 1-figure supplement 2:**
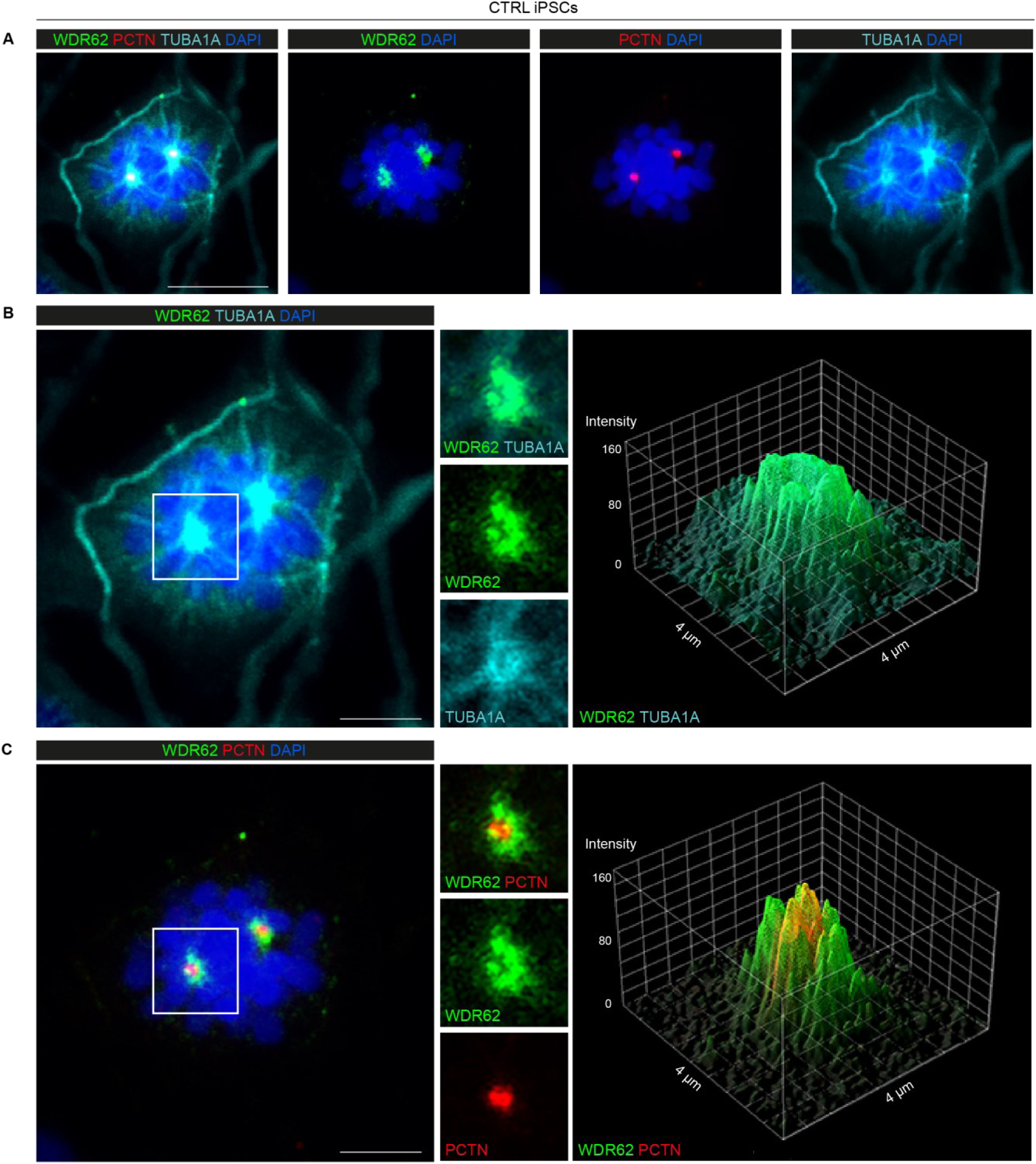
WDR62 is localized at the spindle poles, rather than centrosomes, during mitosis in CTRL iPSCs. A) Representative immunofluorescence assay for WDR62, PCTN, and TUBA1A showing WDR62 signal surrounding centrosomes (PCTN). B) Magnified view of the mitotic cell in A) showing the WDR62 and TUBA1A signals overlapping at the spindle poles (left). The 3D-Surface plot of fluorescence intensity shows a similar WDR62 and TUBA1A signal distribution at the spindle poles (right). C) Magnification of the mitotic cell in A) showing WDR62 surrounding the PCTN signal (left). The 3D-Surface plot of fluorescence intensity shows non-overlapping WDR62 and PCTN signal distribution at the spindle poles (right). Scale bar = 10 μm in A and 5 μm in B, C.

**Figure 2-figure supplement 1:**
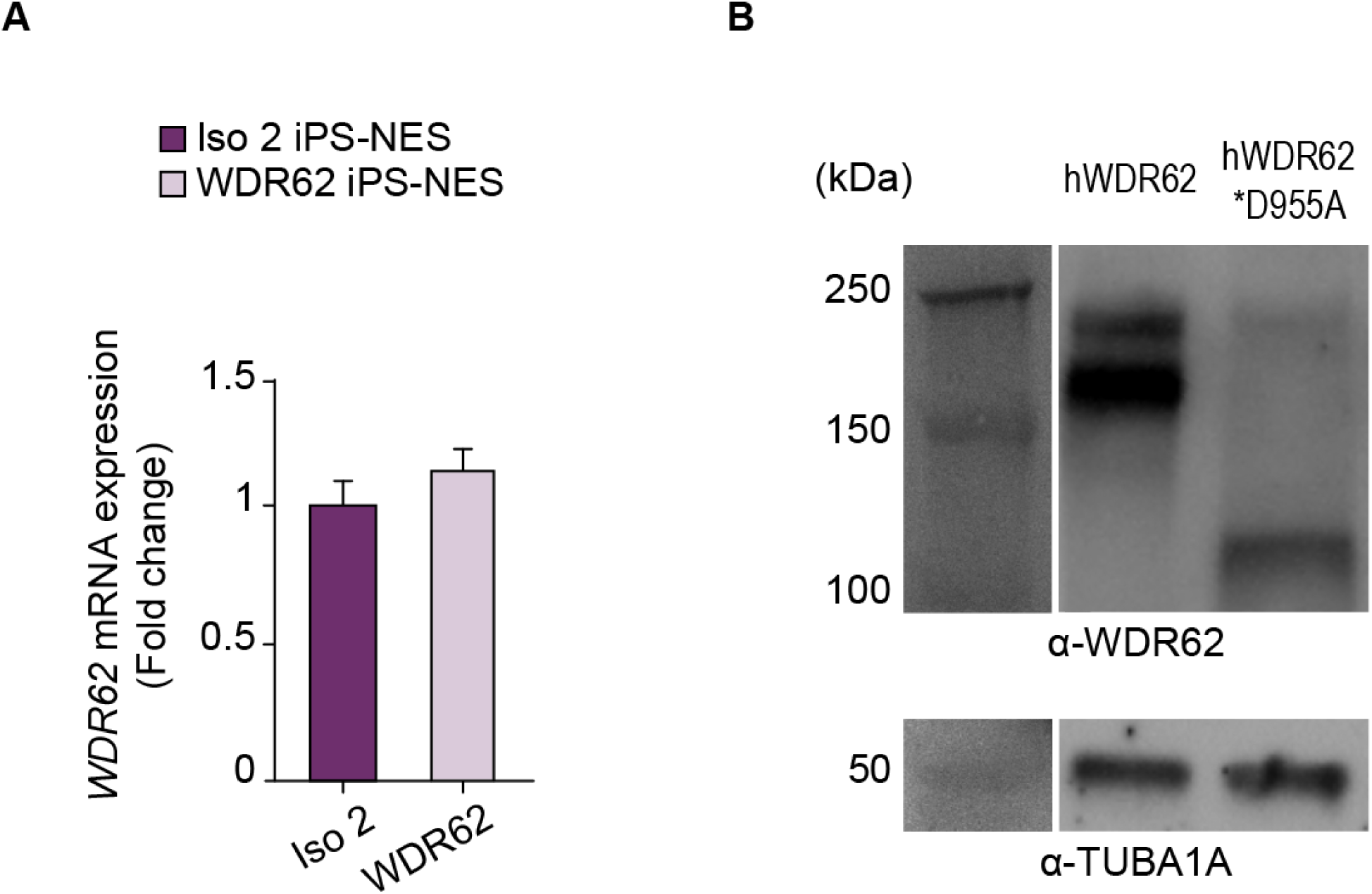
D955AfsX112 mutant WDR62 (D955A) is present in iPS-NES cells. A) RT-qPCR detects *WDR62* mRNA in Iso and WDR62 iPS-NES cells. Quantitative analysis shows no differences in *WDR62* mRNA expression. Data are shown as fold change in *WDR62* gene expression relative to *GAPDH*, a housekeeping gene, according to the 2^−ΔΔCT^ method. B) Western blot showing WDR62 (165 KDa) and mutant WDR62 (118 KDa) protein expression. CTRL iPS-NES cells were transfected with hWDR62 or D955AfsX112 cDNA. Transfected cell lysates were immunoblotted with antibodies against WDR62 and TUBA1A, showing the presence of both WDR62 and mutant WDR62 proteins. **Figure 2-figure supplement 1-source data 1:** Raw data and annotated uncropped western blots from Figure 2-figure supplement 1.

**Figure 3-figure supplement 1:**
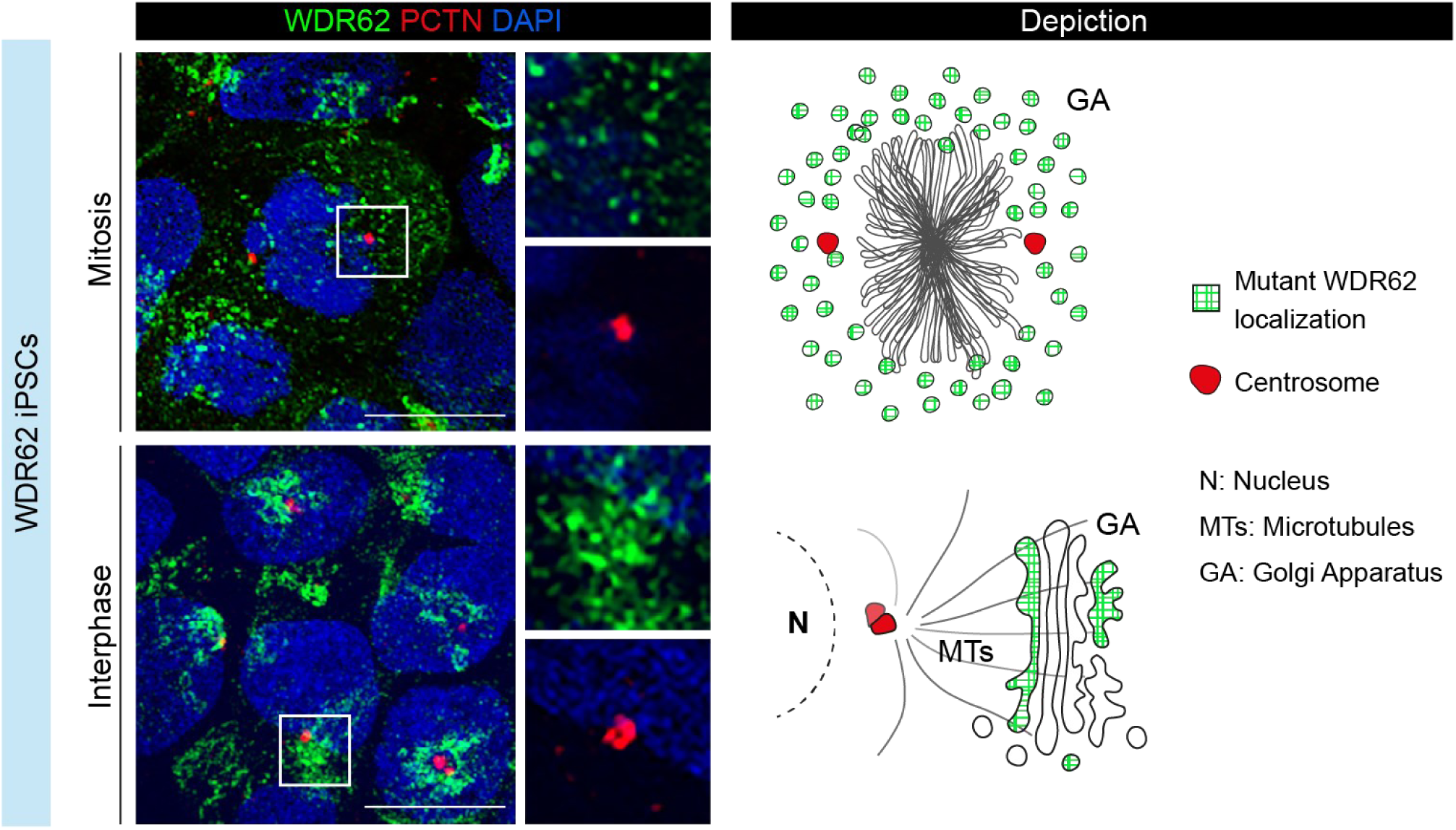
Endogenous WDR62 signal is polarized and perinuclear in WDR62 iPS cells. Representative immunofluorescence assay for WDR62 and PCTN during mitosis and interphase. Magnified view of areas delineated by white squares depicting subcellular localization of mutant WDR62 protein. Schematic representation of mutant WDR62 signal (green) distribution during mitosis (diffuse) and interphase (in Golgi apparatus). Scale bar = 20 μm.

**Figure 3-figure supplement 2:**
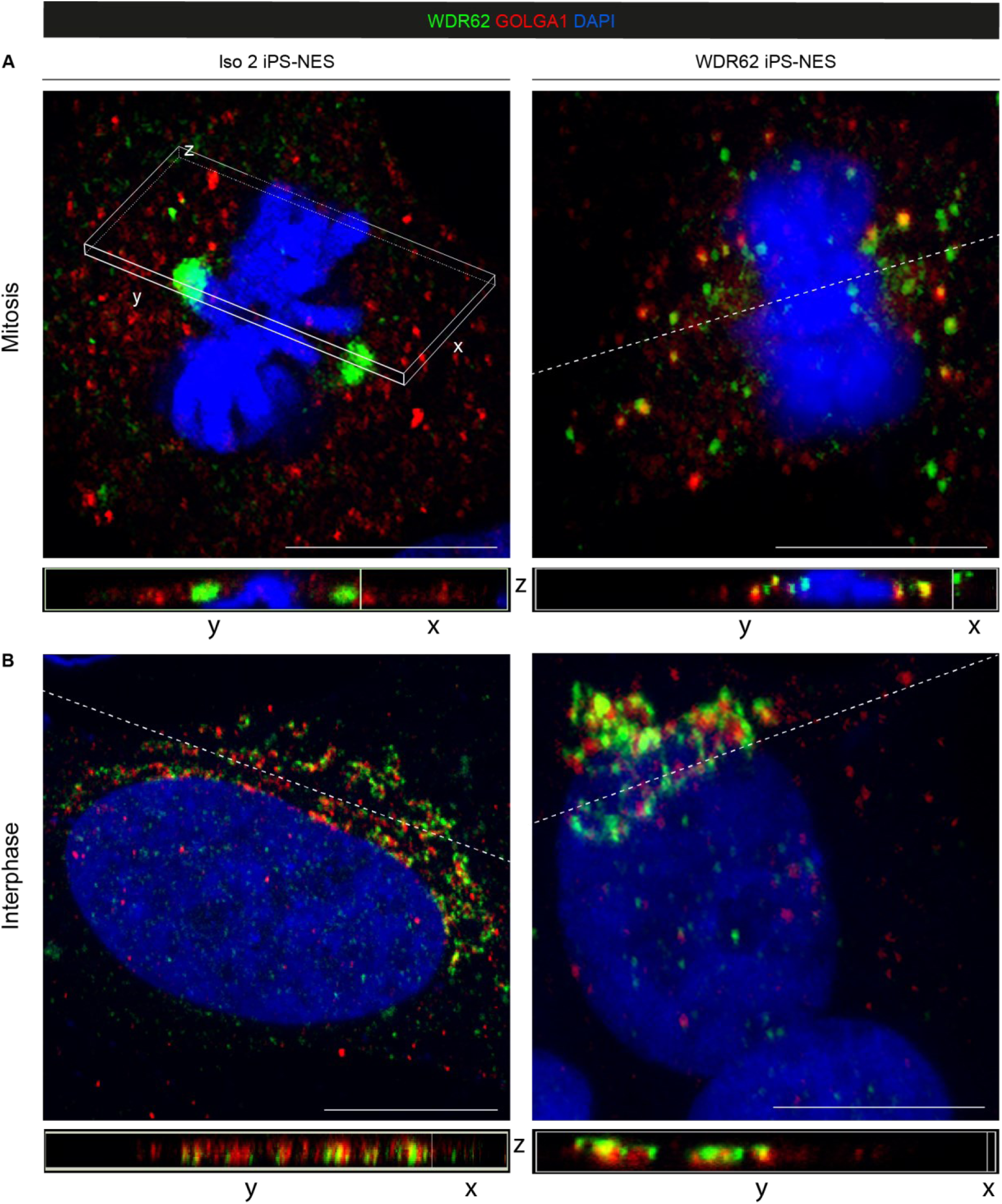
Divergent behaviour of WDR62 in Iso and WDR62 iPS-NES cells. A) High-resolution confocal images and 3D projections showing WDR62 retention to the Golgi apparatus in WDR62 iPS-NES cells during mitosis, contrary to the spindle pole localization in Iso iPS-NES cells. B) During interphase, the WDR62 and Golgi signals overlap as highlighted in the 3D projections. Scale bar = 10 μm in A, B.

**Figure 3-figure supplement 3:**
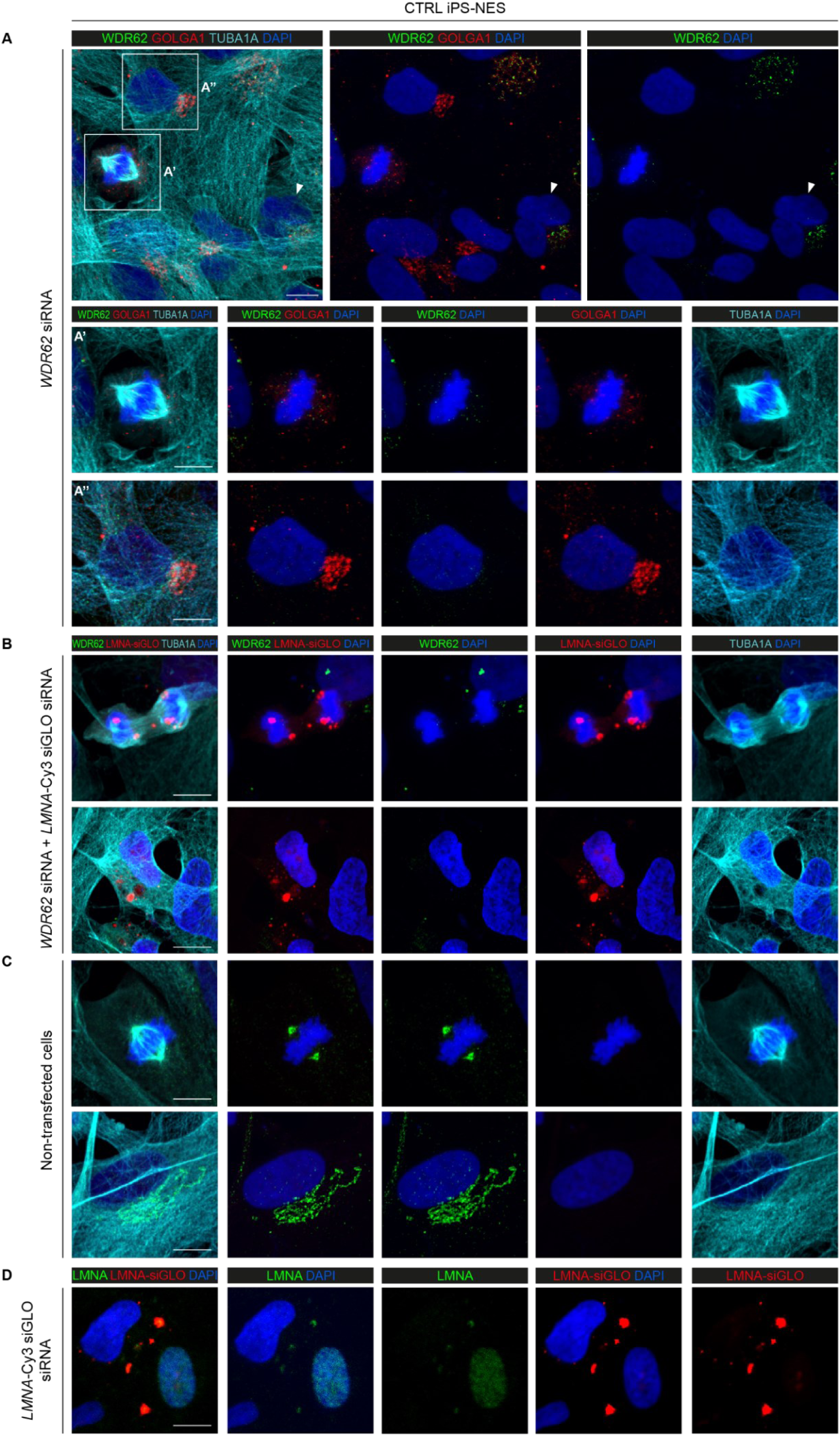
siRNA-mediated knockdown leads to loss of endogenous WDR62 signal. A) Immunofluorescence assay for WDR62, GOLGA1 (also known as Golgin97), and TUBA1A shows no WDR62 signal in CTRL iPS-NES cells transfected with *WDR62*-siRNA. In non-transfected cells (arrowhead), WDR62 signal at the Golgi apparatus is still present. Magnification of *WDR62*-siRNA treated cells in mitosis A**’**) and interphase A**’’**). B) Immunofluorescence assay for WDR62, *LMNA*-Cy3 siGLO siRNA, and TUBA1A in CTRL iPS-NES cells co-transfected with *WDR62*- and *LMNA*-Cy3 siRNAs. *LMNA*-Cy3 siGLO siRNA was used as positive transfection control. In co-transfected cells, WDR62 signal is absent. C) Representative confocal images of WDR62, *LMNA*-Cy3 siGLO siRNA, and TUBA1A in non-transfected CTRL iPS-NES cells, in which WDR62 signal is detected in both mitotic and interphase cells. D) Representative confocal images of LMNA and *LMNA*-Cy3 siGLO siRNA in CTRL iPS-NES cells transfected with *LMNA*-Cy3 siGLO siRNA. Contrary to non-transfected cells, transfected cells show no LMNA signal. Scale bar = 10 μm in A, A’, A’’, B, C, D.

**Figure 3-figure supplement 4:**
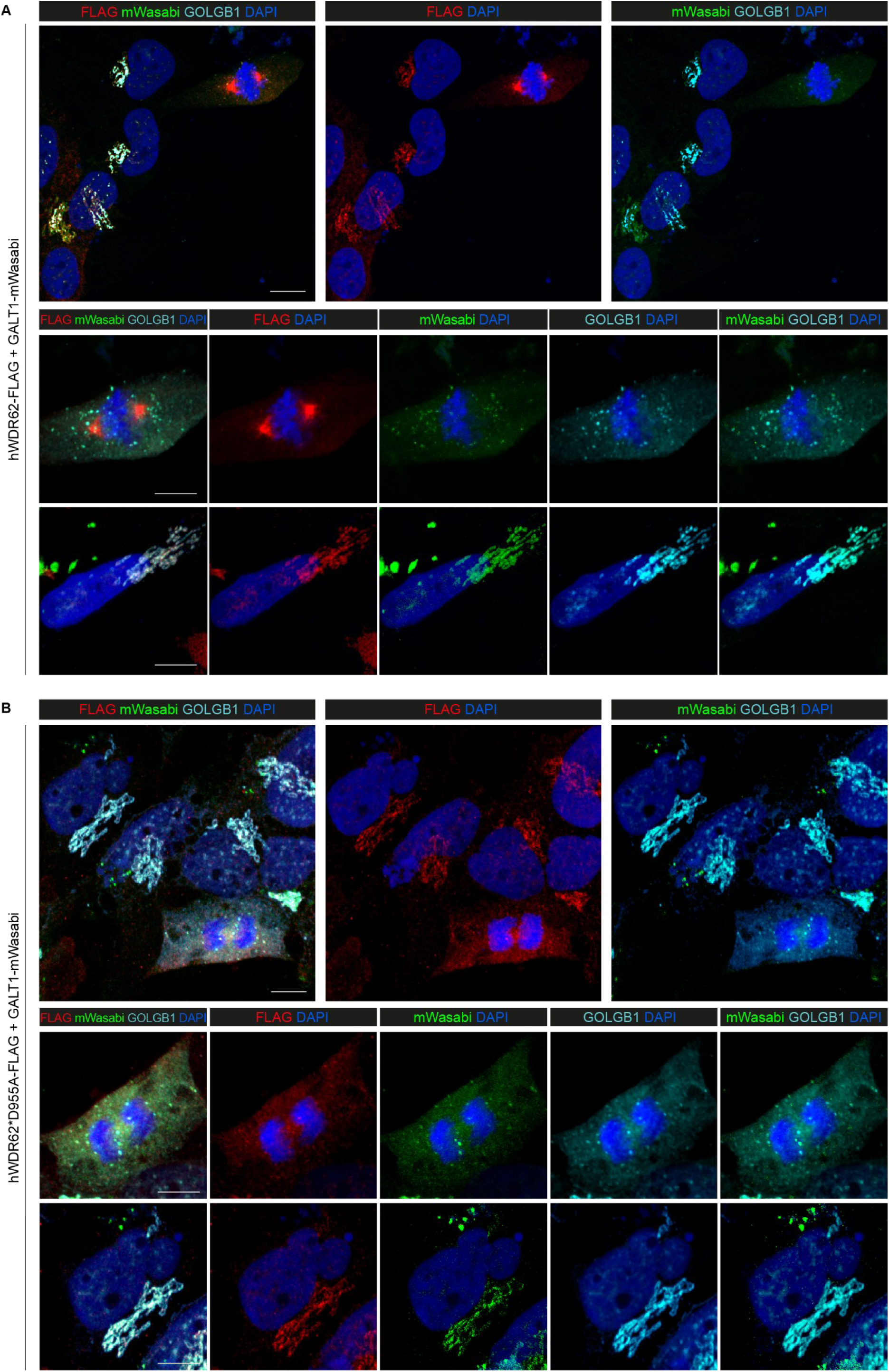
WDR62 localization to the Golgi apparatus during mitosis and interphase. A) CTRL iPS-NES cells were co-transfected with hWDR62-FLAG and GALT1-mWasabi. Confocal imaging for FLAG, mWasabi (a green fluorescent protein) and GOLGB1 (also known as Giantin). Magnification of specific subcellular compartments showing WDR62 localization during mitosis and interphase. B) CTRL iPS-NES cells were co-transfected with hWDR62*D955AfsX112-FLAG and GALT1-mWasabi. Confocal analysis for FLAG, mWasabi, and GOLGB1. Magnification of specific subcellular compartments showing WDR62 localization during mitosis and interphase. The FLAG and mWasabi signals overlap in a manner similar to the endogenous WDR62/Golgi signals. Scale bar = 10 μm in A, B.

**Figure 3-figure supplement 5:**
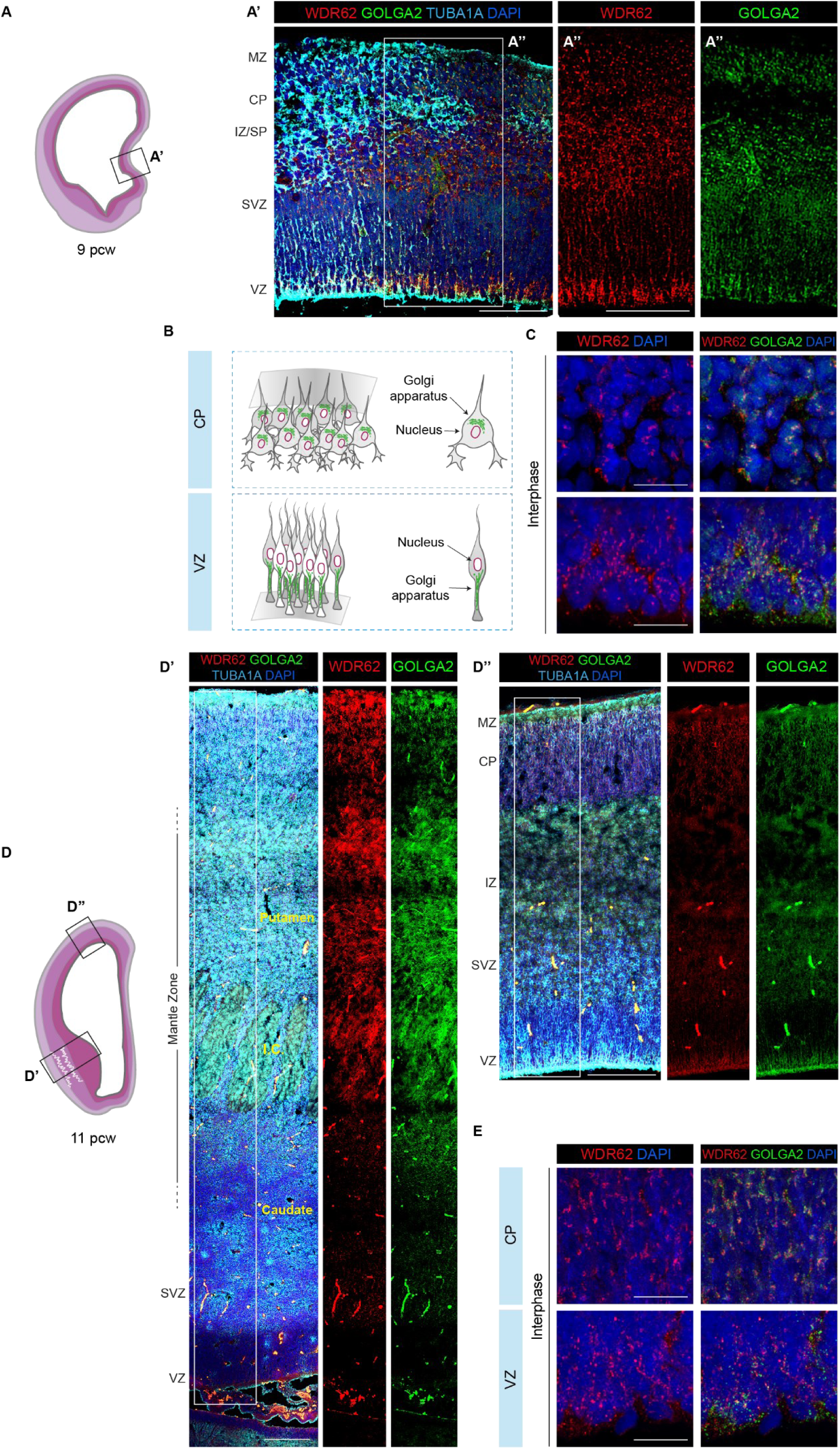
Dynamic expression pattern and subcellular localization of WDR62 in human telencephalon. A) Schematic illustration of a coronal hemisection of human telencephalon at 9 post-conceptional weeks (pcw). A**’**) Representative immunofluorescence assay for WDR62, GOLGA2, and TUBA1A in the VZ of medial pallium. B) Schematic illustration of Golgi apparatus in NPCs and early post-mitotic neurons during cortical development. C) Representative magnified view of WDR62 and GOLGA2 signal distribution during interphase at CP and VZ at 9 pcw. In early post mitotic neurons at the CP, WDR62 shows a discrete perinuclear pattern and follows a diffuse-to-punctate pattern along NPC processes at the VZ, overlapping with GOLGA2. D) Schematic illustration of a coronal hemisection of human telencephalon at 11 pcw. D’, D’’) Representative immunofluorescence assay for WDR62, GOLGA2, and TUBA1A in the developing basal ganglia (D’) and neocortex (D”). E) Representative magnified views of WDR62 and GOLGA2 signal distribution during interphase at the CP and VZ at 11 pcw. WDR62 overlaps with GOLGA2 in post-mitotic neurons at the CP and NPC at the VZ, similar to the pattern observed at 9 pcw. Scale bar = 100 μm in A’, 5 μm in C, E and 100 μm in D’, D’’. MZ (Marginal Zone); CP (Cortical Plate); IZ/SP (Intermediate Zone/Subplate); SVZ (Sub-Ventricular Zone); VZ (Ventricular Zone); I.C. (Internal Capsule).

**Figure 4-figure supplement 1:**
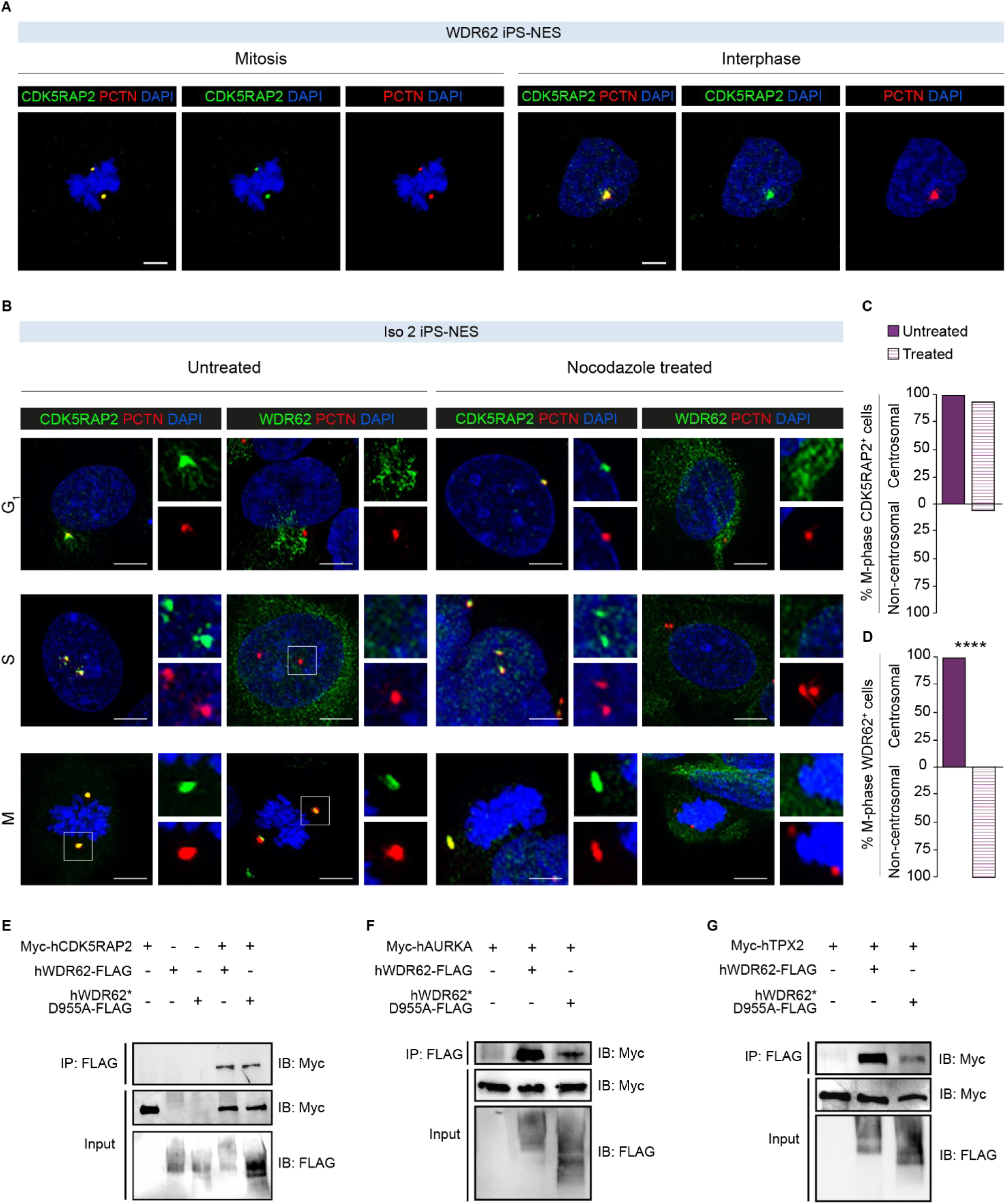
WDR62 interaction with CDK5RAP2, AURKA, and TPX2. A) Immunofluorescence assay for CDK5RAP2 and PCTN in WDR62 iPS-NES cells during mitosis and interphase. B) Representative immunofluorescence assay for CDK5RAP2, WDR62, and PCTN in nocodazole-treated (1.5 μg/ml for 1 h) and untreated Iso iPS-NES cells during G_1_, S, and mitosis (M-phase). C) Quantitative analysis of CDK5RAP2 centrosomal localization during mitosis, which is unaltered by nocodazole. D) Quantitative analysis of WDR62 spindle pole localization during M-phase; nocodazole treatment results in a dispersed perinuclear signal. Data are shown as mean, *p*-value > 0.05 (Chi-square test) and mean, *p*-value < 0.05 (Fisher’s exact test), respectively (total cells, N = 46). E-G) WDR62 interacts with CDK5RAP2 (E), AURKA (F), and TPX2 (G). HEK293T cells were transfected with Myc-hCDK5RAP2, hAURKA, or hTPX2 and hWDR62-FLAG or WDR62 D955AfsX112-FLAG cDNA, as indicated. Cell lysates were immunoprecipitated with an antibody against the Myc-tag and immunoblotted with an antibody against FLAG. WDR62 interaction is detectable with all three partners. Scale bar = 5 μm in A and 10 μm in B. **Figure 4-figure supplement 1-source data 1:** Raw data and annotated uncropped western blots from Figure 4-figure supplement 1. **Figure 4-figure supplement 1-source data 2:** Raw data and annotated uncropped western blots from Figure 4-figure supplement 1. **Figure 4-figure supplement 1-source data 3:** Raw data and annotated uncropped western blots from Figure 4-figure supplement 1.

